# Phylogenetic inference identifies two eumetazoan TRPM clades and an 8^th^ family of TRP channel, TRP soromelastatin (TRPS)

**DOI:** 10.1101/860445

**Authors:** Nathaniel J. Himmel, Thomas R. Gray, Daniel N. Cox

**Affiliations:** Neuroscience Institute, Georgia State University, Atlanta, GA, 30303

## Abstract

TRP melastatins (TRPMs) are most well-known as cold and menthol sensors, but are in fact broadly critical for life, from ion homeostasis to reproduction. Yet the evolutionary relationship between TRPM channels remains largely unresolved, particularly with respect to the placement of several highly divergent members. To characterize the evolution of TRPM and like channels, we performed a large-scale phylogenetic analysis of >1,300 TRPM-like sequences from 14 phyla (Annelida, Arthropoda, Brachiopoda, Chordata, Cnidaria, Echinodermata, Hemichordata, Mollusca, Nematoda, Nemertea, Phoronida, Priapulida, Tardigrada, and Xenacoelomorpha), including sequences from a variety of recently sequenced genomes that fill what would otherwise be substantial taxonomic gaps. These findings suggest: (1) The previously recognized TRPM family is in fact two distinct families, including canonical TRPM channels, and an 8^th^ major, previously undescribed family of animal TRP channel, TRP soromelastatin (TRPS); (2) two TRPM clades predate the last bilaterian-cnidarian ancestor; and (3) the vertebrate-centric trend of categorizing TRPM channels as 1-8 is inappropriate for most phyla, including other chordates.

## Introduction

Transient receptor potential (TRP) channels are a superfamily of ion channel commonly characterized by their 6 transmembrane segments and broad sensory capacity. Among animals, TRP channels have been canonically divided into 7 families (Venkatachalam and Montell 2007; Peng, et al. 2015): TRPA (ankyrin), TRPC (canonical), TRPM (melastatin), TRPML (mucolipin), TRPN (no mechanoreceptor potential C), TRPP (polycystin, or polycystic kidney disease), and TRPV (vanilloid; and a proposed sister family, TRPVL). These TRP channels vary substantially, but TRPM has arguably diversified the most with respect to function, participating in at least cardiac activity (Yue, et al. 2015), magnesium homeostasis (Schlingmann, et al. 2007; Hofmann, et al. 2010), egg activation (Carlson 2019), sperm thermotaxis (De Blas, et al. 2009), cell adhesion (Su, et al. 2006), apoptosis (Driscoll, et al. 2017), inflammation (Ramachandran, et al. 2013), and most famously, cold (Bautista, et al. 2007; Turner, et al. 2016) and menthol (McKemy, et al. 2002; Peier, et al. 2002; Himmel, et al. 2019) sensing.

TRPM channels are also thought to be incredibly ancient, predating the emergence of metazoans (>1000Ma) (Peng, et al. 2015; Himmel, et al. 2019). In some species, single moonlighting proteins carry out several functions (*e.g. Drosophila melanogaster* Trpm), while in others, functions are compartmentalized in a set of diverse paralogues (*e.g.* human TRPM1-8). However, little is known about the evolutionary history of TRPMs, or to what degree channels are related across taxa. Our understanding of TRPM evolution is additionally clouded by the existence of several highly divergent putative TRPM channels with uncertain origins (Teramoto, et al. 2005; Peng, et al. 2015; Kozma, et al. 2018; Himmel, et al. 2019). Here, we made use of the rapidly growing body of genomic data in order to better characterize the evolution of the TRPM family.

Via a stringent screening process, we assembled a database of >1,300 predicted TRPM-like sequences from 14 diverse eumetazoan phyla (**Fig. 1**). In this database we gave particular attention to underrepresented taxa, as well as included TRP genes identified in a number of recently sequenced genomes (**Table S1**; including, but not limited to, acoel flatworm, moon jelly, and great white shark). Herein, we elucidate the evolutionary history and familial organization of both TRP melastatin, and a previously unrecognized sister family that predates the Cnidaria-Bilateria split, TRP soromelastatin.

**Fig. 1.**
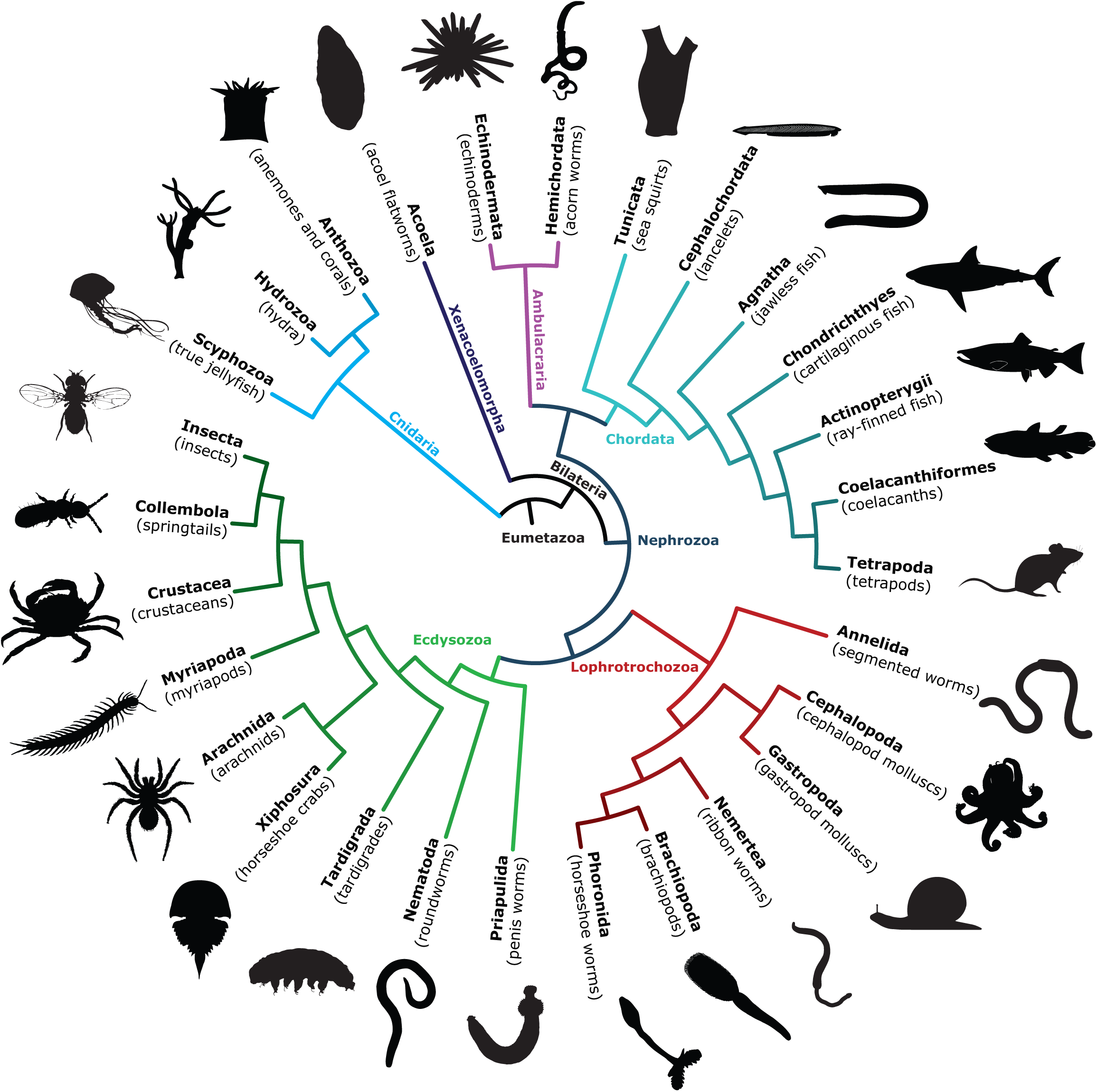
Major, widely-recognized taxa included in these phylogenetic analyses, and relationships assumed throughout. TRPM and TRPM-like sequences were collected for 318 species, from 14 diverse eumetazoan phyla (Annelida, Arthropoda, Brachiopoda, Chordata, Cnidaria, Echinodermata, Hemichordata, Mollusca, Nematoda, Nemertea, Phoronida, Priapulida, Tardigrada, and Xenacoelomorpha).

## Materials & Methods

### Data Collection & Curation

Starting with previously characterized TRPM sequences from human (NCBI CCDS), mouse (NCBI CCDS), *Drosophila melanogaster* (FlyBase), and *Caenorhabditis elegans* (WormBase), a TRPM-like protein sequence database was assembled by performing BLASTp against NCBI collections of non-redundant protein sequences, with *D. melanogaster* Trpm (isoform RE, FlyBase ID: FBtr0339077) serving as the bait sequence. In order to maximize useful phylogenetic information, only BLAST hits >300 amino acids in length with an E-value less than 1E-30 were retained. As we were interested in the origins of TRPM channels, and in less-studied taxa, only three tetrapod sequence-sets were included, from human, mouse, and chicken.In order to expand the taxa sampled, tBLASTn and BLASTp were used to search publically available, genomically-informed gene models for 11 cnidarians, 2 xenacoelomorphs, 1 hemichordate, 1 nemertean, 1 phoronid, 2 agnathans, and 4 chondrichthyes (**Table S1**).

We used several methods in order to validate and improve the quality of the initial database. First, CD-HIT (threshold 90% similarity) was used to identify and remove duplicate sequences and predicted isoforms, retaining the longest isoform (Li and Godzik 2006; Huang, et al. 2010; Fu, et al. 2012). Phobius was then used to predict transmembrane topology (Käll, et al. 2004, 2007); sequences which did not have at least 6 predicted transmembrane (TM) segments, which is typical of TRP channels, were removed. Sequences with more than the 6 predicted TM segments were analyzed via InterProScan (Mitchell, et al. 2019), and those with more than 1 ion-transport domain were removed. More than 90% of the remaining sequences contained a highly conserved glycine residue in the predicted TM domain (corresponding to *D. melanogaster* G-1049); the vast majority of those missing this residue had large gaps in an initial alignment and were subsequently removed.

Searches for TRPS (ced-11-like), TRPN, and TRPC sequences followed the same protocol. For TRPS, sequences from *Caenorhabditis elegans, Strigamia maritima*, and *Octopus vulgaris* were used as bait. For TRPN and TRPC datasets, *Drosophila melanogaster* nompC (isoform PA, FlyBase ID: FBpp0084879) and Trp (isoform PA, Flybase ID: FBpp0084879) served as bait sequences, respectively.

### Principal Component Analysis

Principal component analysis (PCA) was used to help resolve protein families. TRPC, TRPN, and TRPM/TRPS database sequences were aligned by MAFFT. A pairwise sequence identity matrix was then computed, and PCA performed against it in Jalview (Waterhouse, et al. 2009). Data were exported from Jalview and visualized and edited in GraphPad Prism and Adobe Illustrator CS6.

### Phylogenetic Tree Estimation

For the maximum likelihood approach, sequences were first aligned using MAFFT with default settings (Rozewicki, et al. 2019). Gap rich sites and poorly-aligned sequences were trimmed with TrimAl (Capella-Gutiérrez, et al. 2009). IQ-Tree (Nguyen, et al. 2014) was then used to generate trees by the maximum likelihood approach, using the best models automatically selected by ModelFinder (Kalyaanamoorthy, et al. 2017). Branch support was calculated by ultrafast bootstrapping (UFBoot, 2000 bootstraps) (Hoang, et al. 2017).

In order to test the alternative hypothesis that some trees formed due to long-branch attraction, gs2 was used to generate trees by the Graph Splitting method (Matsui and Iwasaki 2019). Branch support values were computed by the packaged edge perturbation method (EP, 2000 iterations). All trees were visualized and edited in iTOL and Adobe Illustrator CS6.

### Homologue Prediction via Tree Reconciliation

In order to identify duplication events, TRPS and TRPM phylograms were reconciled using NOTUNG 2.9.1 (Durand, et al. 2006; Vernot, et al. 2008; Stolzer, et al. 2012). Edge weight threshold was set to 1.0, and the costs of duplications and losses were set to 1.5 and 1.0, respectively. In order to formulate the most parsimonious interpretation of the resulting trees, weak branches were rearranged (UFboot 95 cutoff) against a cladogram based in an NCBI taxonomic tree, wherein we placed Xenacoelomorpha (represented by acoel flatworms) as the sister group to all other bilaterians (Cannon, et al. 2016), and Priapulida as an outgroup to all other ecdysozoans (Yamasaki, et al. 2015). All other polytomies were randomly resolved using the ape package in R (Paradis and Schliep 2018).

## Results

### An ancient, unrecognized sister family to TRPM – TRP soromelastatin (TRPS)

Proteins within the same family typically have a high degree of sequence similarity, yet highly divergent TRPM-like proteins have been catalogued, a notable example being *Caenorhabditis elegans* cell death abnormal 11 (ced-11). The canonical *C. elegans* TRPMs gtl-1, gtl-2, and gon-2 share roughly 40% sequence identity with each other. However, ced-11—often considered a fourth *C. elegans* TRPM—shares approximately 18% sequence identity with the 3 canonical paralogues. Given this substantial difference, it seemed plausible that ced-11, and like proteins, had been errantly included in the TRPM family.

The TRPC and TRPN families are typically thought to be most closely related to TRPM (Peng, et al. 2015), and therefore constituted hypothetical homes for ced-11. ced-11, however, shares only 15% sequence identity with known *C. elegans* TRPC paralogues (trp-1 and trp-2), and 14% sequence identity with *C. elegans* TRPN (trp-4). Yet sequence identity between trp-4 and its TRPC counterparts is approximately 20%. In other words, ced-11 is less similar to TRPMs than TRPNs and TRPCs are to each other.

In order to clarify the relationship of ced-11-like proteins to canonical TRPM channels, we collected those sequences most similar to it from our initial TRPM-like sequence database, and phylogenetically characterized them. BLASTing our database with ced-11-like sequences recovered a number of sequences restricted to several protostome taxa and lancelets (Cephalochordata).

For any species with a ced-11-like protein, we assembled a database of putative TRPC and TRPN channel sequences. These sequences were then phylogenetically characterized alongside cnidarian, xenacoelomorph, insect (*D. melanogaster*), and human sequences. In the resulting tree, ced-11-like proteins formed a sister clade to the more traditional TRPM clade, the latter including cnidarian TRPM-like channels (**Fig. 2** and **Fig. S1**).

**Fig. 2.**
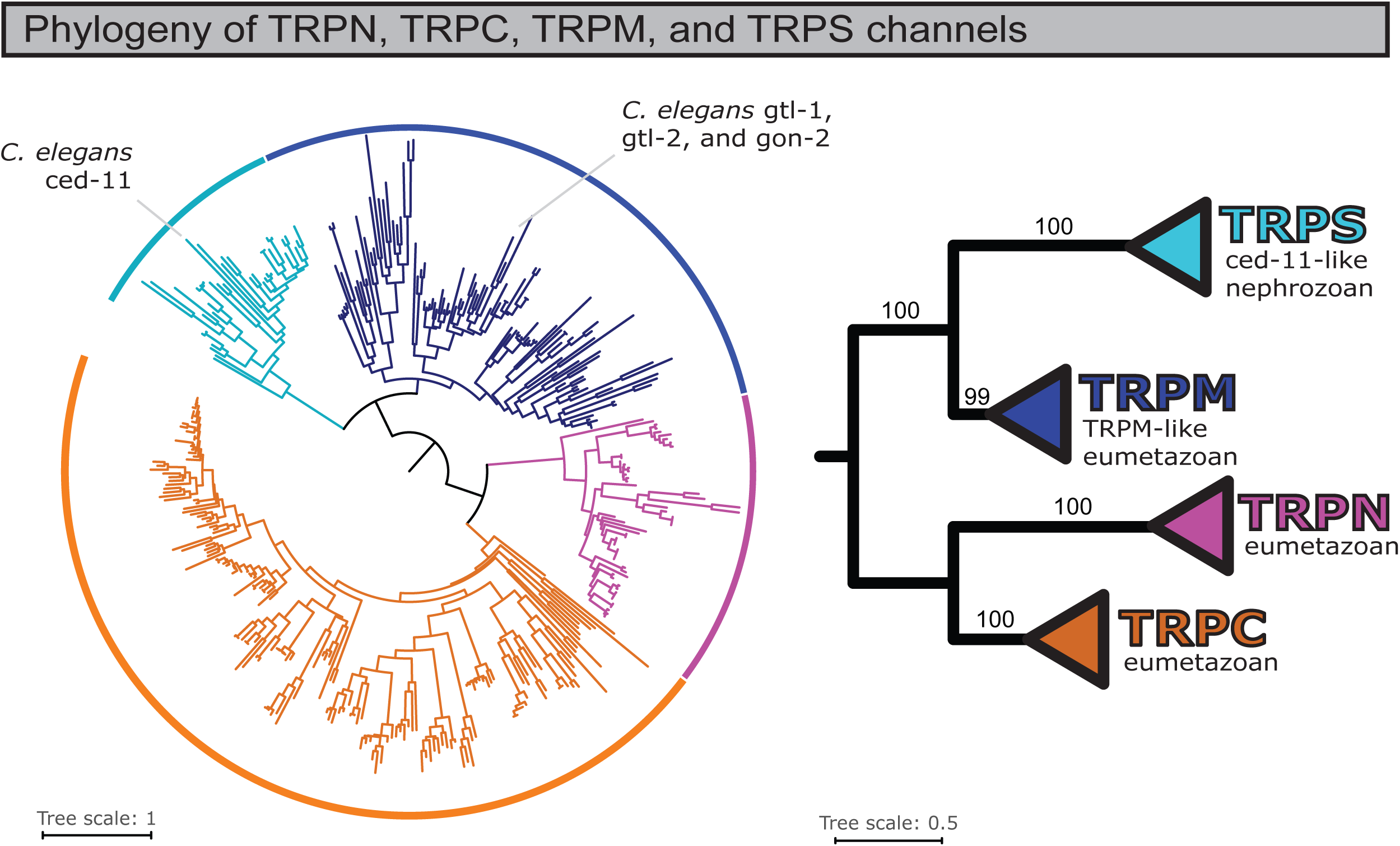
*Caenorhabditis elegans* ced-11, and ced-11-like sequences, belong to a previously unrecognized family of TRP channels, the TRP soromelastatins (TRPS). TRPM-like and ced-11-like sequences form two distinct clades, with the topology suggesting divergence prior to the Cnidaria-Bilateria split. Left, maximum likelihood tree showing the relationship between traditional TRPM and TRPS/ced-11-like sequences among those species that have TRPS/ced-11-like species. Right, summary and branch support (UFboot) for indicated clades.

Two competing hypotheses could explain these findings: (1) ced-11-like proteins constitute a distinct family of TRP channel which predates the cnidarian-bilaterian split, or (2) a variety of TRPM channels emerged independently in various taxa and diversified extremely rapidly, resulting in a clade which formed as a result of long-branch attraction, an artifact of many phylogenetic analyses (Bergsten 2005).

Hypothesis 2 appears highly unlikely. Most importantly, while *C. elegans* ced-11 itself has a relatively long branch, when qualitatively compared to other clades, the branches within the ced-11-like clade were not unusually long (**Fig. 2 and Fig. S1**). Additionally, principal component analysis of a pairwise sequence identity matrix revealed that ced-11-like sequences cluster together independent of TRPM-like sequences, suggesting they cluster in the phylogram due to sequence similarity (**Fig. 3A**). We tested the long-branch hypothesis by estimating trees which excluded Cnidaria and Xenacoelomorpha, which had particularly long branches and could serve to exacerbate long-branch attraction, were it present. The resultant phylogram still evidenced the split between ced-11-like and TRPM-like channels, with high branch confidence (**Fig. S2**). Moreover, we generated a phylogram by the Graph Splitting method, which is reported to be extremely robust when faced with the possibility of long-branch attraction in superfamily-level datasets (Matsui and Iwasaki 2019). This method likewise reproduced the ced-11-like-TRPM split with high edge perturbation branch support (**Fig. S3**).

**Fig. 3.**
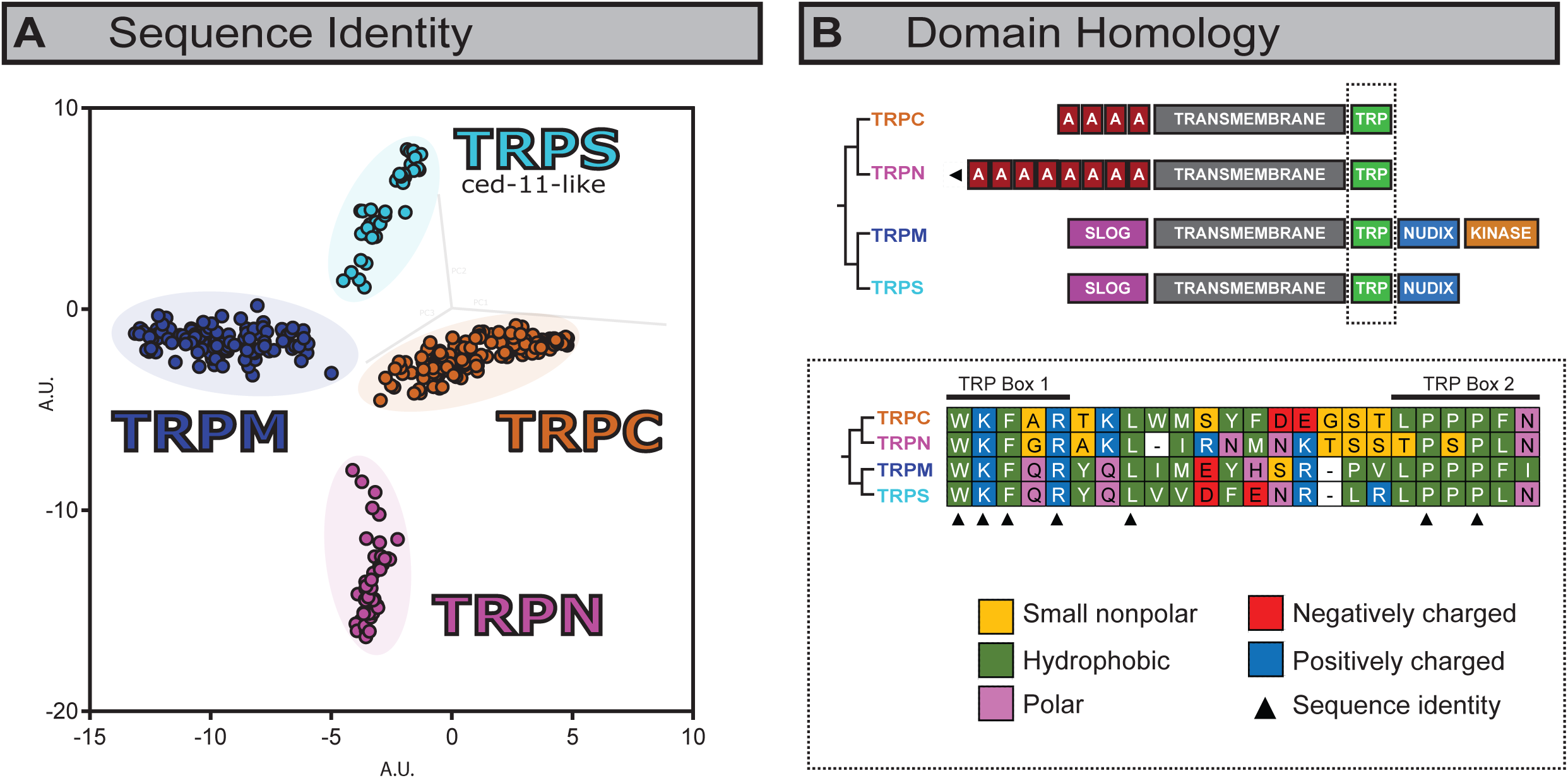
The predicted structures of putative TRPS channels suggest a SLOG- and Nudix-linked, Ankyrin-free TRPS-TRPM ancestor. (**A**) Principal component analyses of pairwise sequence identity for alignment of TRPC, TRPN, TRPM, and ced-11-like (TRPS) sequences show 4 distinct clusters. 3-dimensional PCA plot extracted from Jalview and plotted in two transformed dimensions. (**B**) Like TRPM channels, TRPS channels have both SLOG and Nudix domains, but lack the kinase domain associated with some TRPM channels. Moreover, these TRPS channels have a divergent consensus sequence in the highly conserved TRP domain.

These results strongly indicate that these two lineages diverged in or prior to the last cnidarian-bilaterian ancestor, and that ced-11-like proteins constitute an 8^th^ family of metazoan TRP channel. We have thus named the ced-11-like family of TRP channels TRP soromelastatin (*soro-*, sister), or TRPS (**Fig. 2** and **Fig. 3**).

### The structure of TRPS channels suggests a SLOG- and Nudix-linked ancestor

While the function of TRPS channels remains unknown, domain prediction reveals that both TRPM and TRPS channels share an N-terminal SMF/DprA-LOG (SLOG) domain (which is hypothesized to function in ligand sensing) and a C-terminal ADP-ribose phosphohydrolase (Nudix) domain (**Fig. 3B**, top). These results indicate that the ancestral TRPM-TRPS channel was likely both SLOG- and Nudix-linked, and that the Ankyrin repeats typical of TRPC and TRPNs were lost prior to the TRPM-TRPS split. The TRPM alpha kinase domain (typical of human TRPM6 and TRPM7), however, appears to have arisen specifically in the TRPM lineage. These findings are consistent with previous findings suggesting that the TRPM ancestor was Nudix-linked (Schnitzler, et al. 2008), but pushes the origins of this domain further back in evolutionary history.

A notable difference between TRPM and TRPS channels lies in the TRP domain, a highly conserved, hydrophobic region located C-terminally to the transmembrane domain of TRPC, TRPN, and TRPM channels (Venkatachalam and Montell 2007). Consensus sequences for TRPM and TRPS (**Fig. 2**), while identical in TRP box 1, are divergent in TRP box 2 and the intermediate TRP segment (**Fig. 3B**, bottom). The functional consequences of these changes, if any, are unknown.

### The TRPS family is largely restricted to protostomes

Having established that these TRPS sequences constitute a distinct set of channels, we assembled a more complete TRPS sequence database and phylogenetically characterized the channel family. These data suggest that, among Eumetazoa, TRPS genes are only present in some protostomes and lancelets (**Fig. 4** and **Fig. S4**). The lack of widespread conservation among deuterostomes (most notably vertebrates) and insects likely explains why the family had gone unnoticed until now.

**Fig. 4.**
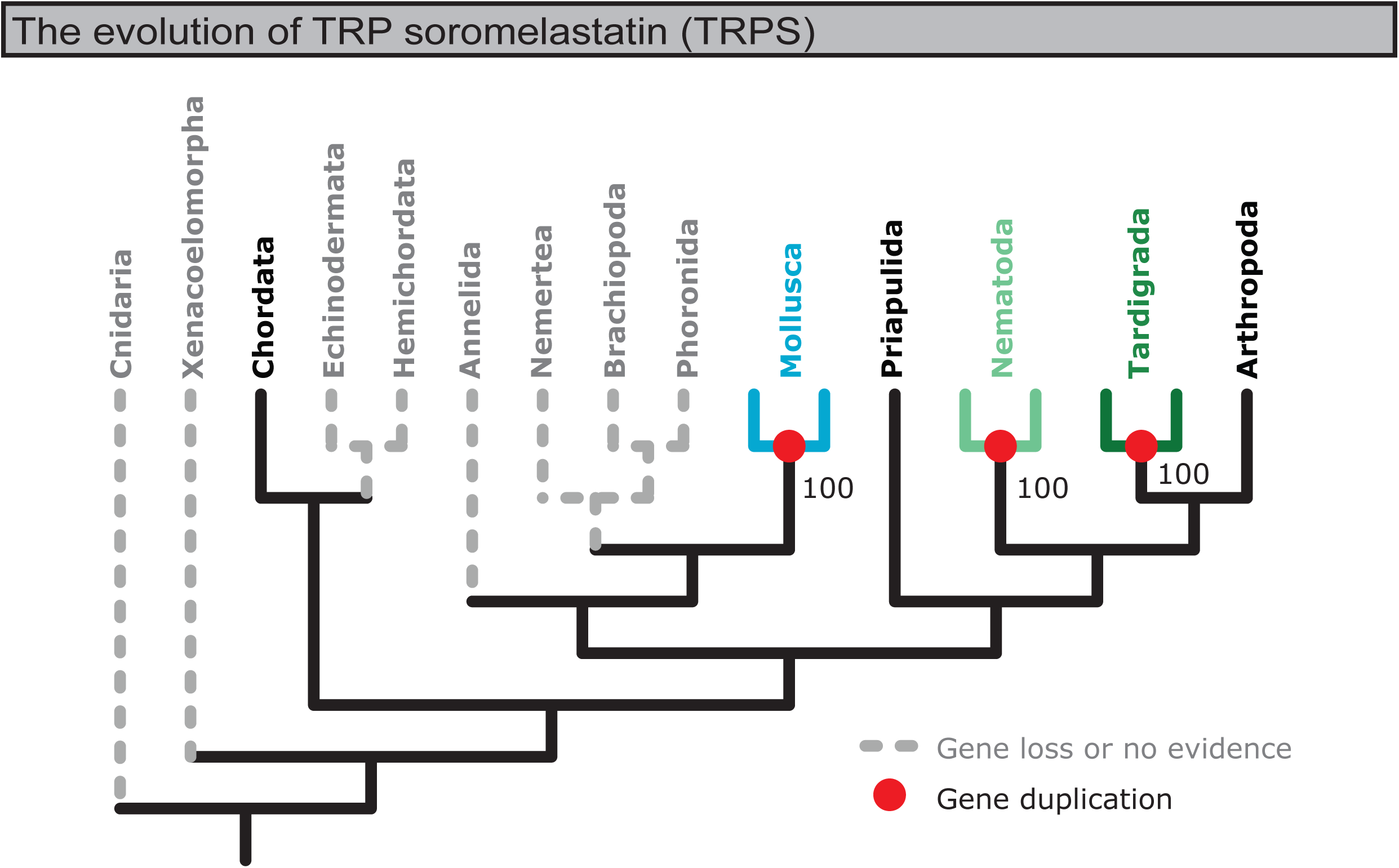
The evolution of TRPS channels. The TRPS family is largely restricted to protostome lineages. Figure derived from reconciled maximum likelihood tree of TRPS sequences (**Fig. S3**). Red dots and colored branches indicate phylum-specific duplication events. Grey and dashed branches indicate that no TRPS sequences were found for the indicated phylum, and were inferred to be loss events.

TRPS was likely lost early in deuterostome evolution – among the ambulacrarians (echinoderms and hemichordates), and in early Olfactores (tunicates and vertebrates) following the olfactore-lancelet split (**Fig. S5**, left). A recent study evidences that Ambulacraria and Xenacoelomorpha might form sister clades (Philippe, et al. 2019) – if this is the case, it may be more likely that TRPS was lost early in so-called “xenambulacrarian” evolution (**Fig. S5**, right).

TRPS duplication appears to have been limited during early animal evolution. While the number of TRPS paralogues varies by species (**Fig. S4**), duplication events occurred only after major taxa emerged, independently in molluscs, nematodes, tardigrades, and chelicerates (including arachnids and horseshoe crabs). Molluscs have two TRPS paralogues, but present lack of evidence for lophotrochozoan TRPS outside of molluscs makes it difficult to predict at what point in spiralian evolution the duplication event occurred. The simplest explanation is that it occurred specifically in molluscs, and that a single TRPS copy was lost among other lophotrochozoan taxa.

Among Euarthropoda, TRPS appears in chelicerates and myriapods, but there is no evidence for TRPS in crustaceans, springtails, or insects, suggesting that the single arthropod TRPS was lost in Pancrustacea, conserved in Myriapoda, and expanded independently in Chelicerata (**Fig. S6**).

### Two TRPM clades predate the Cnidaria-Bilateria split

We next phylogenetically characterized and reconciled TRPM sequences among major taxa. Each set of sequences was initially assessed alongside cnidarian, xenacoelomorph, *Drosophila*, and human sequences, and rooted with TRPS sequences.

The general consensus of these analyses indicates that the TRPM family is made up of two distinct clades, here and previously deemed αTRPM and βTRPM (Himmel, et al. 2019), which emerged prior to the Cnidaria-Bilateria split (**Fig. 5** and **Figs. S7-S16**). What might have constituted a previously described basal clade can be almost wholly explained by the discovery of TRPS (Peng, et al. 2015; Himmel, et al. 2019). In some of our initial phylograms, a basal or separate clade did appear, yet it always included Xenacoelomorpha and was inconsistent in its topology across analyses (**Figs. S7-S11**), suggesting that Xenacoelomorpha acted as a phylogenetically unstable rogue taxon (Thomson and Shaffer 2010). In order to assess this possibility, we performed a second set of analyses which excluded Xenacoelomorpha. This resulted in trees with largely consistent topology despite differing taxon sampling, indicating that xenacoelomorph sequences are in fact problematic (**Figs. S12-S16**).

**Fig. 5.**
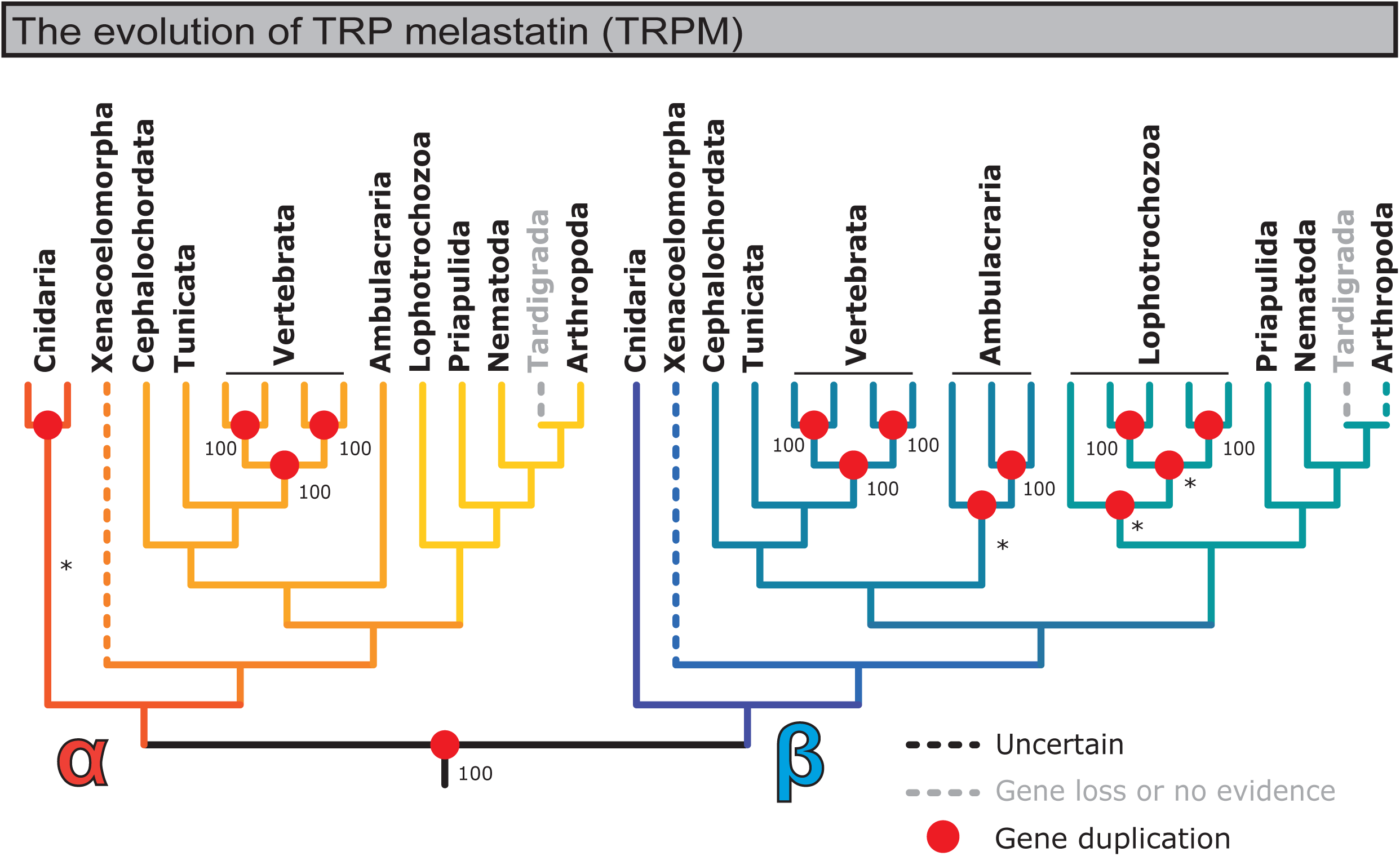
Summary of the evolution of TRPM channels. The TRPM family is widely conserved, and is present in all phyla surveyed except Tardigrada. Figure derived from consensus topologies in trees generated against TRPM database sequences, with branch support for duplication branches extracted from phylograms without Xenacoelomorpha (**Fig. S12-S16**). Asterisk (*) indicates that a duplication branch most frequently formed due to rearrangement (initial UFboot branch support <95). Here, phyla were expanded/collapsed to more easily show TRPM diversification: Chordata was expanded into Cephalochordata (lancelets), Tunicata, and Vertebrata; Echinodermata and Hemichordata were collapsed into Ambulacraria; and Annelida, Nemertea, Brachiopoda, Phoronida, and Mollusca were collapsed into Lophotrochozoa. Red dots indicate duplication events. Dashed lines in color indicate sequences considered *incertae sedis*. Dashed lines in grey indicate that no sequences were found for the indicated taxon, and were inferred to be loss events.

Due to the overwhelming consistency of trees with different taxon sampling, and the inconsistency seen in trees including Xenacoelomorpha, xenacoelomorph TRPM sequences were treated as rogue taxa. In addition, an extremely small subset of arthropod TRPMs (12 sequences restricted to chelicerates and crustaceans; **Fig. S11** and **Fig. S16**) may be part of a previously described Crustacea-specific TRPM sub-family (Kozma, et al. 2018). These trees suggest that these sequences are βTRPM-like and related to a subset of cnidarian sequences, yet this Cnidaria-inclusive clade is not strongly evidenced in phylograms with different taxon sampling. Like Xenacoelomorph sequences, the evolutionary histories of these sequences are left *incertae sedis*.

In summary, these results strongly support two duplication events predating the Cnidaria-Bilateria split: the TRPS-TRPM split and the α-β TRPM split.

### TRPM1-8 expansion occurred early in vertebrate evolution, and constitutes a poor standard for TRPM family organization

The vertebrate TRPM1-8 expansion has been the focus of the majority of TRPM literature, and has been the principal basis for characterizing TRPM channels (Samanta, et al. 2018; Zhang, et al. 2018; Chen, et al. 2019). However, these trees evidence that the TRPM1-8 expansion occurred after the vertebrate-tunicate split, and before agnathans (jawless fish; lampreys and hagfish) split from the ancestor of all other vertebrates (**Fig. 5, Fig. 6**, and **Fig. S13**). Although immunohistochemical evidence has previously suggested that TRPM8 is present in teleost fish (Majhi, et al. 2015), we found no evidence of it in available sequences for ray-finned fish, cartilaginous fish, or agnathans (**Fig. 6** and **Fig. S17**). While the simplest naive hypothesis would be that TRPM8 did not emerge until lobe-finned fish emerged, these phylogenetic analyses indicate that TRPM8 was independently lost in the indicated taxa, and conserved in the lobe-finned vertebrate lineage (including tetrapods).

**Fig. 6.**
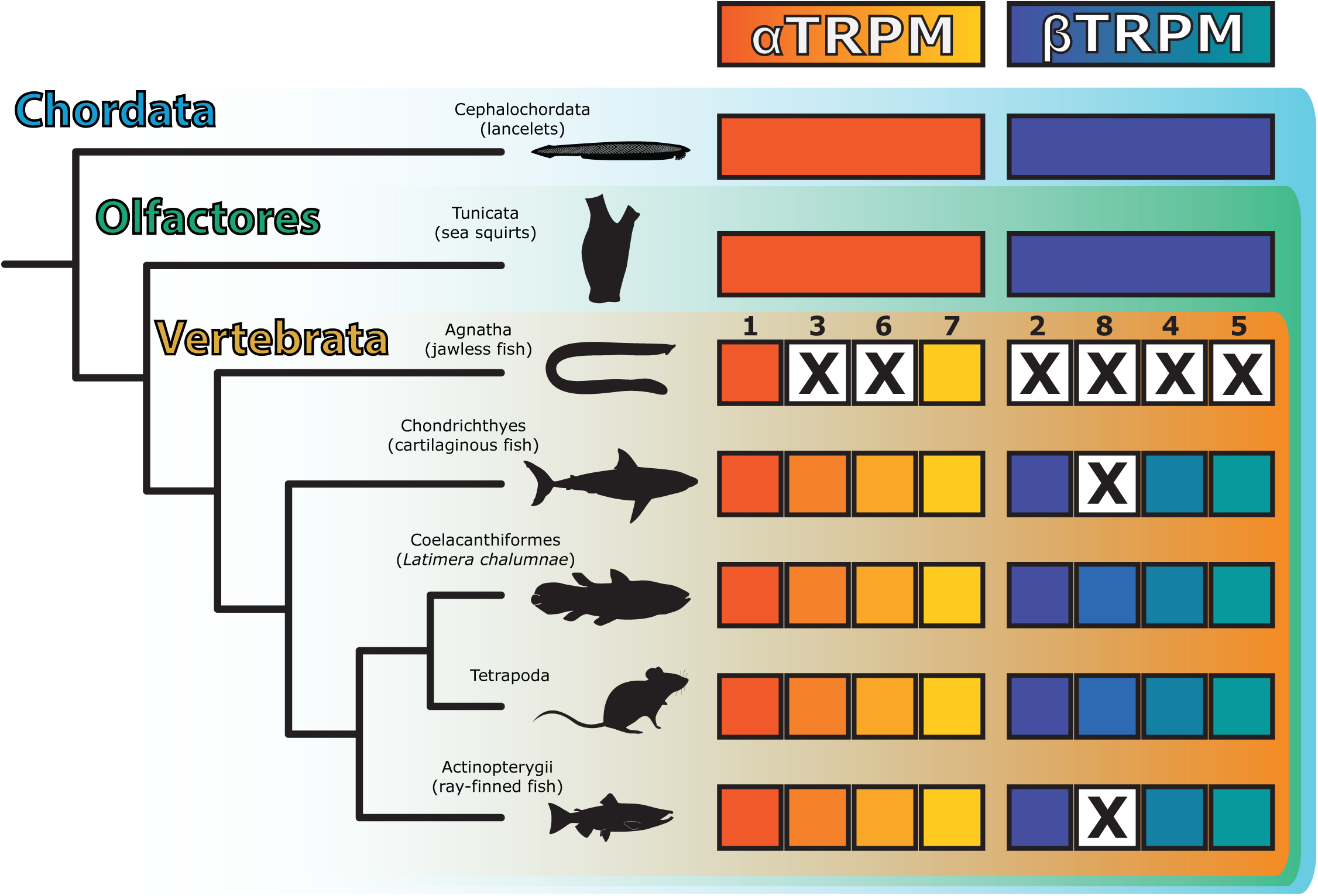
TRPM1-8 diversified soon after the vertebrate-tunicate split, and there is no evidence for TRPM8 in jawless, cartilaginous, or ray-finned fish. Figure derived from phylogram of chordate TRPM sequences (**Fig. S17**). An “X” indicates inferred gene loss, or lack of evidence of that gene in the indicated taxon.

Moreover, the 1-8 nomenclature may under-describe TRPMs among one of the most abundant vertebrate clades – the teleost fish. While basal ray-finned fish (e.g., *Erpetoichthys calabaricus*, the freshwater snakefish, or reedfish) have a TRPM topology that closely matches other vertebrates, the emergence of teleosts came with TRPM expansion. For example, there are as many as three teleost TRPM4 paralogues (**Fig. S17**).

## Discussion

The evolutionary history of TRPM channels has been clouded by divergent sequences, making it uncertain if an ancestral clade of TRPMs had survived in species like *C. elegans*, or if these species had independently evolved rapidly changing TRPM paralogues (Teramoto, et al. 2005; Peng, et al. 2015; Kozma, et al. 2018; Himmel, et al. 2019). By taking advantage of the abundance of publicly available genomic data, we have demonstrated that the difficulty in phylogenetically characterizing TRPM channels is the result of an ancient, hidden family of channels that appeared before the Cnidaria-Bilateria split – the TRPS. By recognizing and characterizing this family, we now better understand not only the evolution and diversification of TRPM, but also the evolution of the broader TRP superfamily.

While some have been careful in describing TRP channels in taxon-specific ways (Saito and Shingai 2006; Hofmann, et al. 2010; Peng, et al. 2015), these findings are the strongest challenge to the pervasive, vertebrate-centric dogma that the TRPM family is constituted by 8 distinct paralogues organized into four subfamilies (Samanta, et al. 2018; Zhang, et al. 2018; Chen, et al. 2019). These results instead support that the eumetazoan TRPM family consists of two distinct radiations (αTRPM and βTRPM) which themselves predate the Cnidaria-Bilateria split. Importantly, these findings support that TRPM diversification occurred independently among cnidarians, ambulacrarians, lophotrochozoans, and other taxa, and that the TRPM1-8 expansion is specific to vertebrates. Based on these findings, we conclude that the TRPM1-8 nomenclature is at best evolutionarily uninformative (*e.g.* insect channels being simply TRPM1- or 3-like), and at worst grossly inaccurate (*e.g.* cnidarian TRPMs belonging to the TRPM2/8 subfamily) for describing members of this diverse family of critically important ion channels.

## Acknowledgments

We thank Dr. Charles Derby and Mihika Kozma for critically assessing the original manuscript, and the PhyloPic repository, the source for many of the animal silhouettes used throughout (distributed in public domain). We also thank all the investigators who made sequence information public, which made this work possible.

## Author contributions

Conceptualization and methodology, NJH; sequence collection, NJH; database curation, NJH and TRG; domain homology analysis, NJH and TRG; phylogenetic and other formal analyses, NJH; prepared the original draft, NJH; reviewed and edited the final draft, NJH, TRG, and DNC; visualization, NJH; supervision, DNC; funding acquisition, DNC.

## Data and materials availability

The TRPN, TRPC, TRPM, and TRPS sequence databases have been deposited on Dryad in the FASTA format (doi:10.5061/dryad.kwh70rz03).

## References

Arimoto A, Hikosaka-Katayama T, Hikosaka A, Tagawa K, Inoue T, Ueki T, Yoshida M-a, Kanda M, Shoguchi E, Hisata K, et al. 2019. A draft nuclear-genome assembly of the acoel flatworm *Praesagittifera naikaiensis*. GigaScience 8.

Baumgarten S, Simakov O, Esherick LY, Liew YJ, Lehnert EM, Michell CT, Li Y, Hambleton EA, Guse A, Oates ME, et al. 2015. The genome of *Aiptasia*, a sea anemone model for coral symbiosis. Proceedings of the National Academy of Sciences 112:11893.

Bautista DM, Siemens J, Glazer JM, Tsuruda PR, Basbaum AI, Stucky CL, Jordt S-E, Julius D. 2007. The menthol receptor TRPM8 is the principal detector of environmental cold. Nature 448:204–208.

Bergsten J. 2005. A review of long-branch attraction. Cladistics 21:163–193.

Cannon JT, Vellutini BC, Smith J, Ronquist F, Jondelius U, Hejnol A. 2016. Xenacoelomorpha is the sister group to Nephrozoa. Nature 530:89.

Capella-Gutiérrez S, Silla-Martínez JM, Gabaldón T. 2009. trimAl: a tool for automated alignment trimming in large-scale phylogenetic analyses. Bioinformatics (Oxford, England) 25:1972–1973.

Carlson AE. 2019. Mechanical stimulation activates *Drosophila* eggs via Trpm channels. Proceedings of the National Academy of Sciences 116:18757.

Chen Y, Zhang X, Yang T, Bi R, Huang Z, Ding H, Li J, Zhang J. 2019. Emerging structural biology of TRPM subfamily channels. Cell Calcium 79:75–79.

Cunning R, Bay RA, Gillette P, Baker AC, Traylor-Knowles N. 2018. Comparative analysis of the *Pocillopora damicornis* genome highlights role of immune system in coral evolution. Scientific Reports 8:16134.

De Blas GA, Darszon A, Ocampo AY, Serrano CJ, Castellano LE, Hernández-González EO, Chirinos M, Larrea F, Beltrán C, Treviño CL. 2009. TRPM8, a versatile channel in human sperm. PLOS ONE 4:e6095–e6095.

Driscoll K, Stanfield GM, Droste R, Horvitz HR. 2017. Presumptive TRP channel CED-11 promotes cell volume decrease and facilitates degradation of apoptotic cells in *Caenorhabditis elegans*. Proceedings of the National Academy of Sciences 114:8806–8811.

Durand D, Halldórsson BV, Vernot B. 2006. A Hybrid Micro–Macroevolutionary Approach to Gene Tree Reconstruction. Journal of Computational Biology 13:320–335.

Fu L, Niu B, Zhu Z, Wu S, Li W. 2012. CD-HIT: accelerated for clustering the next-generation sequencing data. Bioinformatics (Oxford, England) 28:3150–3152.

Gehrke AR, Neverett E, Luo Y-J, Brandt A, Ricci L, Hulett RE, Gompers A, Ruby JG, Rokhsar DS, Reddien PW, et al. 2019. Acoel genome reveals the regulatory landscape of whole-body regeneration. Science 363:eaau6173.

Gold DA, Katsuki T, Li Y, Yan X, Regulski M, Ibberson D, Holstein T, Steele RE, Jacobs DK, Greenspan RJ. 2019. The genome of the jellyfish *Aurelia* and the evolution of animal complexity. Nature Ecology & Evolution 3:96–104.

Hara Y, Yamaguchi K, Onimaru K, Kadota M, Koyanagi M, Keeley SD, Tatsumi K, Tanaka K, Motone F, Kageyama Y, et al. 2018. Shark genomes provide insights into elasmobranch evolution and the origin of vertebrates. Nature Ecology & Evolution 2:1761–1771.

Himmel NJ, Letcher JM, Sakurai A, Gray TR, Benson MN, Cox DN. 2019. *Drosophila* menthol sensitivity and the Precambrian origins of transient receptor potential-dependent chemosensation. Philosophical Transactions of the Royal Society B: Biological Sciences 374:20190369.

Hoang DT, Chernomor O, von Haeseler A, Minh BQ, Vinh LS. 2017. UFBoot2: Improving the Ultrafast Bootstrap Approximation. Molecular Biology and Evolution 35:518–522.

Hofmann T, Chubanov V, Chen X, Dietz AS, Gudermann T, Montell C. 2010. *Drosophila* TRPM Channel Is Essential for the Control of Extracellular Magnesium Levels. PLOS ONE 5:e10519.

Huang Y, Niu B, Gao Y, Fu L, Li W. 2010. CD-HIT Suite: a web server for clustering and comparing biological sequences. Bioinformatics (Oxford, England) 26:680–682.

Käll L, Krogh A, Sonnhammer ELL. 2007. Advantages of combined transmembrane topology and signal peptide prediction—the Phobius web server. Nucleic Acids Research 35:W429–W432.

Käll L, Krogh A, Sonnhammer ELL. 2004. A Combined Transmembrane Topology and Signal Peptide Prediction Method. Journal of Molecular Biology 338:1027–1036.

Kalyaanamoorthy S, Minh BQ, Wong TKF, von Haeseler A, Jermiin LS. 2017. ModelFinder: fast model selection for accurate phylogenetic estimates. Nature Methods 14:587.

Kozma MT, Schmidt M, Ngo-Vu H, Sparks SD, Senatore A, Derby CD. 2018. Chemoreceptor proteins in the Caribbean spiny lobster, *Panulirus argus*: Expression of Ionotropic Receptors, Gustatory Receptors, and TRP channels in two chemosensory organs and brain. PLOS ONE 13:e0203935.

Li W, Godzik A. 2006. Cd-hit: a fast program for clustering and comparing large sets of protein or nucleotide sequences. Bioinformatics 22:1658–1659.

Luo Y-J, Kanda M, Koyanagi R, Hisata K, Akiyama T, Sakamoto H, Sakamoto T, Satoh N. 2018. Nemertean and phoronid genomes reveal lophotrochozoan evolution and the origin of bilaterian heads. Nature Ecology & Evolution 2:141–151.

Majhi RK, Saha S, Kumar A, Ghosh A, Swain N, Goswami L, Mohapatra P, Maity A, Kumar Sahoo V, Kumar A, et al. 2015. Expression of temperature-sensitive ion channel TRPM8 in sperm cells correlates with vertebrate evolution. PeerJ 3:e1310–e1310.

Marra NJ, Stanhope MJ, Jue NK, Wang M, Sun Q, Pavinski Bitar P, Richards VP, Komissarov A, Rayko M, Kliver S, et al. 2019. White shark genome reveals ancient elasmobranch adaptations associated with wound healing and the maintenance of genome stability. Proceedings of the National Academy of Sciences 116:4446.

Matsui M, Iwasaki W. 2019. Graph Splitting: A Graph-Based Approach for Superfamily-Scale Phylogenetic Tree Reconstruction. Systematic Biology.

McKemy DD, Neuhausser WM, Julius D. 2002. Identification of a cold receptor reveals a general role for TRP channels in thermosensation. Nature 416:52–58.

Mitchell AL, Attwood TK, Babbitt PC, Blum M, Bork P, Bridge A, Brown SD, Chang H-Y, El-Gebali S, Fraser MI, et al. 2019. InterPro in 2019: improving coverage, classification and access to protein sequence annotations. Nucleic Acids Research 47:D351–D360.

Nguyen L-T, Schmidt HA, von Haeseler A, Minh BQ. 2014. IQ-TREE: A Fast and Effective Stochastic Algorithm for Estimating Maximum-Likelihood Phylogenies. Molecular Biology and Evolution 32:268–274.

Paradis E, Schliep K. 2018. ape 5.0: an environment for modern phylogenetics and evolutionary analyses in R. Bioinformatics 35:526–528.

Peier AM, Moqrich A, Hergarden AC, Reeve AJ, Andersson DA, Story GM, Earley TJ, Dragoni I, McIntyre P, Bevan S, et al. 2002. A TRP Channel that Senses Cold Stimuli and Menthol. Cell 108:705–715.

Peng G, Shi X, Kadowaki T. 2015. Evolution of TRP channels inferred by their classification in diverse animal species. Molecular Phylogenetics and Evolution 84:145–157.

Philippe H, Poustka AJ, Chiodin M, Hoff KJ, Dessimoz C, Tomiczek B, Schiffer PH, Müller S, Domman D, Horn M, et al. 2019. Mitigating Anticipated Effects of Systematic Errors Supports Sister-Group Relationship between Xenacoelomorpha and Ambulacraria. Current Biology 29:1818-1826.e1816.

Ramachandran R, Hyun E, Zhao L, Lapointe TK, Chapman K, Hirota CL, Ghosh S, McKemy DD, Vergnolle N, Beck PL, et al. 2013. TRPM8 activation attenuates inflammatory responses in mouse models of colitis. Proceedings of the National Academy of Sciences 110:7476.

Rozewicki J, Li S, Amada KM, Standley DM, Katoh K. 2019. MAFFT-DASH: integrated protein sequence and structural alignment. Nucleic Acids Research 47:W5–W10.

Saito S, Shingai R. 2006. Evolution of thermoTRP ion channel homologs in vertebrates. Physiological Genomics 27:219–230.

Samanta A, Hughes TET, Moiseenkova-Bell VY. 2018. Transient Receptor Potential (TRP) Channels. Sub-cellular biochemistry 87:141–165.

Schlingmann KP, Waldegger S, Konrad M, Chubanov V, Gudermann T. 2007. TRPM6 and TRPM7—Gatekeepers of human magnesium metabolism. Biochimica et Biophysica Acta (BBA) - Molecular Basis of Disease 1772:813–821.

Schnitzler MMy, Wäring J, Gudermann T, Chubanov V. 2008. Evolutionary determinants of divergent calcium selectivity of TRPM channels. The FASEB Journal 22:1540–1551.

Shinzato C, Shoguchi E, Kawashima T, Hamada M, Hisata K, Tanaka M, Fujie M, Fujiwara M, Koyanagi R, Ikuta T, et al. 2011. Using the *Acropora digitifera* genome to understand coral responses to environmental change. Nature 476:320.

Simakov O, Kawashima T, Marlétaz F, Jenkins J, Koyanagi R, Mitros T, Hisata K, Bredeson J, Shoguchi E, Gyoja F, et al. 2015. Hemichordate genomes and deuterostome origins. Nature 527:459–465.

Smith JJ, Kuraku S, Holt C, Sauka-Spengler T, Jiang N, Campbell MS, Yandell MD, Manousaki T, Meyer A, Bloom OE, et al. 2013. Sequencing of the sea lamprey (*Petromyzon marinus*) genome provides insights into vertebrate evolution. Nature Genetics 45:415.

Stolzer M, Lai H, Xu M, Sathaye D, Vernot B, Durand D. 2012. Inferring duplications, losses, transfers and incomplete lineage sorting with nonbinary species trees. Bioinformatics (Oxford, England) 28:i409–i415.

Su L-T, Agapito MA, Li M, Simonson WTN, Huttenlocher A, Habas R, Yue L, Runnels LW. 2006. TRPM7 regulates cell adhesion by controlling the calcium-dependent protease calpain. The Journal of biological chemistry 281:11260–11270.

Teramoto T, Lambie EJ, Iwasaki K. 2005. Differential regulation of TRPM channels governs electrolyte homeostasis in the C. elegans intestine. Cell metabolism 1:343–354.

Thomson RC, Shaffer HB. 2010. Sparse Supermatrices for Phylogenetic Inference: Taxonomy, Alignment, Rogue Taxa, and the Phylogeny of Living Turtles. Systematic Biology 59:42–58.

Turner HN, Armengol K, Patel AA, Himmel NJ, Sullivan L, Iyer SC, Bhattacharya S, Iyer EPR, Landry C, Galko MJ, et al. 2016. The TRP Channels Pkd2, NompC, and Trpm Act in Cold-Sensing Neurons to Mediate Unique Aversive Behaviors to Noxious Cold in *Drosophila*. Current biology 26:3116–3128.

Venkatachalam K, Montell C. 2007. TRP channels. Annual review of biochemistry 76:387-417.

Vernot B, Stolzer M, Goldman A, Durand D. 2008. Reconciliation with non-binary species trees. Journal of computational biology 15:981–1006.

Voolstra C, Miller D, Ragan M, Hoffmann A, Hoegh-Guldberg O, Bourne D, Ball E, Ying H, Foret S, Takahashi S, et al. 2015. The ReFuGe 2020 Consortium—using “omics” approaches to explore the adaptability and resilience of coral holobionts to environmental change. Frontiers in Marine Science 2.

Voolstra CR, Li Y, Liew YJ, Baumgarten S, Zoccola D, Flot J-F, Tambutté S, Allemand D, Aranda M. 2017. Comparative analysis of the genomes *of Stylophora pistillata* and *Acropora digitifera* provides evidence for extensive differences between species of corals. Scientific Reports 7:17583.

Wang X, Liew YJ, Li Y, Zoccola D, Tambutte S, Aranda M. 2017. Draft genomes of the corallimorpharians *Amplexidiscus fenestrafer* and *Discosoma* sp. Molecular Ecology Resources 17:e187–e195.

Waterhouse AM, Procter JB, Martin DMA, Clamp M, Barton GJ. 2009. Jalview Version 2--a multiple sequence alignment editor and analysis workbench. Bioinformatics (Oxford, England) 25:1189–1191.

Yamasaki H, Fujimoto S, Miyazaki K. 2015. Phylogenetic position of Loricifera inferred from nearly complete 18S and 28S rRNA gene sequences. Zoological letters 1:18–18.

Yue Z, Xie J, Yu AS, Stock J, Du J, Yue L. 2015. Role of TRP channels in the cardiovascular system. American journal of physiology -- Heart and circulatory physiology 308:H157–H182.

Zhang Z, Tóth B, Szollosi A, Chen J, Csanády L. 2018. Structure of a TRPM2 channel in complex with Ca^2+^ explains unique gating regulation. eLife 7:e36409.

