## Supplementary Materials for "Phylogenetic inference identifies two eumetazoan TRPM clades and an 8^th^ family of TRP channel, TRP soromelastatin (TRPS)"

**Title:** Preprint supplement to: **“**Phylogenetic inference identifies two eumetazoan TRPM clades and an 8th family of TRP channel, TRP soromelastatin (TRPS)”

**Authors and Affiliations:**

Nathaniel J. Himmel[1,*], Thomas R. Gray[1], and Daniel N. Cox[1,*]

1 – Neuroscience Institute, Georgia State University, Atlanta, GA, 30303
* - Corresponding authors

**Authors for Correspondence:**

**Nathaniel J. Himmel**
Neuroscience Institute
Georgia State University
P.O. Box 5030
Atlanta, GA 30302-5030


**Daniel N. Cox**
Neuroscience Institute
Georgia State University
P.O. Box 5030
Atlanta, GA 30302-5030


### Supplementary Materials:

| Phylum | Species | Source |
|---|---|---|
| Cnidaria | *Aurelia* (moon jelly) | (Gold, et al. 2019) |
| | *Acropora digitifera* (coral) | (Shinzato, et al. 2011) |
| | *Acropora tenuis* (coral) | (Voolstra, et al. 2015) |
| | *Aiptasia* (sea anemone) | (Baumgarten, et al. 2015) |
| | *Amplexidiscus fenestrafer* (elephant ear anemone) | (Wang, et al. 2017) |
| | *Fungia* spp. (coral) | (Voolstra, et al. 2015) |
| | *Galaxea fascicularis* (galaxy coral) | (Voolstra, et al. 2015) |
| | *Goniastrea aspera* (stony coral) | (Voolstra, et al. 2015) |
| | *Pocillopora damicornis* (lace coral) | (Cunning, et al. 2018) |
| | *Porites lutea* (small polyp stony coral) | (Voolstra, et al. 2015) |
| | *Stylophora pistillata* (hood coral) | (Voolstra, et al. 2017) |
| Xenacoelomorpha | *Hofstenia miamia* (three-banded panther worm) | (Gehrke, et al. 2019) |
| | *Praesagittifera naikaiensis* (acoel flatworm) | (Arimoto, et al. 2019) |
| Hemichordata | *Ptychodera flava* | (Simakov, et al. 2015) |
| Chordata | *Petromyzon marinus* (sea lamprey) | (Smith, et al. 2013) |
| | *Eptatretus burger* (inshore hagfish) | PRJEB21290 |
| | *Chiloscyllium punctatum* (brownband bamboo shark) | (Hara, et al. 2018) |
| | *Rhincodon typus* (whale shark) | (Hara, et al. 2018) |
| | *Scyliorhinus torazame* (cloudy catshark) | (Hara, et al. 2018) |
| | *Carcharodon carcharias* (white shark) | (Marra, et al. 2019) |
| Nemertea | *Notospermus geniculatus* (ribbon worm) | (Luo, et al. 2018) |
| Phoronida | *Phoronis australis* (horseshoe worm) | (Luo, et al. 2018) |

**Table S1.** Genomes added to the initial NCBI-based sequence database.

#### Supplemental Figures

**Fig. S1.** TRPS constitutes a distinct family of TRP channel. Maximum likelihood tree for TRPM, TRPS (ced-11-like), TRPN, and TRPC sequences, for all those species in initial database that had a ced-11-like sequence. UFboot confidence is indicated by red-green color scale, with major branch values listed.

**Fig. S2.** Maximum likelihood tree for TRPM, TRPS (ced-11-like), TRPN, and TRPC sequences for all those species in initial database that had a ced-11-like sequence, but excluding Xenacoelomorpha and Cnidaria in order to test for effects of long-branch attraction. UFboot confidence is indicated by red-green color scale, with major branch values listed.

**Fig. S3.** Graph Splitting tree for TRPM, TRPS (ced-11-like), TRPN, and TRPC sequences for all those species in initial database that had a ced-11-like sequence, in order to test for effects of long-branch attraction. Edge perturbation (EP) confidence is indicated by red-green color scale, with major branch values listed.

**Fig. S4.** Reconciled and rearranged maximum likelihood TRPS phylogram with duplication sites (red) and UFboot branch support values listed. Branches without support values were rearranged (<95 UFboot) by NOTUNG. Individual expansion events occurred in molluscs, nematodes, tardigrades, and chelicerates. While *S. maritima* has 2 TRPS genes, this was not assumed to represent a taxon-wide duplication event due to it being the sole representative of Myriapoda.

**Fig. S5.** Two hypotheses concerning the loss of TRPS in Ambulacraria and Olfactores. While a monophyletic Deuterostomia has been well supported for some time (left), recent work has suggested that Ambulacraria may be a sister clade to Xenacoelomorpha (Philippe, et al. 2019) (right).

**Fig. S6.** Arthropod TRPS diversification, extracted from **Fig. S3**. The duplication in arthropod TRPS appears restricted to Chelicerata, and the simplest hypothesis concerning TRPS loss is that it was lost early in the evolution of Pancrustacea.

**Fig. S7.** Reconciled and rearranged maximum likelihood tree of TRPM sequences from Ambulacraria (bold), with TRPM sequences from Cnidaria, Xenacoelomorpha, human, and *Drosophila* for context, and TRPS sequences for rooting.

**Fig. S8.** Reconciled and rearranged maximum likelihood tree of TRPM sequences from Chordata (bold, excluding ray-finned fish), with TRPM sequences from Cnidaria, Xenacoelomorpha, human, and *Drosophila* for context, and TRPS sequences for rooting.

**Fig. S9.** Reconciled and rearranged maximum likelihood tree of TRPM sequences from Lophotrochozoa (bold), with TRPM sequences from Cnidaria, Xenacoelomorpha, human, and *Drosophila* for context, and TRPS sequences for rooting.

**Fig. S10.** Reconciled and rearranged maximum likelihood tree of TRPM sequences from Priapulida and Nematoda (bold), with TRPM sequences from Cnidaria, Xenacoelomorpha, human, and *Drosophila* for context, and TRPS sequences for rooting.

**Fig. S11.** Reconciled and rearranged maximum likelihood tree of TRPM sequences from Priapulida and Arthropoda (bold), with TRPM sequences from Cnidaria, Xenacoelomorpha, human, and *Drosophila* for context, and TRPS sequences for rooting.

**Fig. S12.** Reconciled and rearranged maximum likelihood tree of TRPM sequences from Ambulacraria (bold) as in **Fig. S7**, with Xenacoelomorpha removed. Ambulacrarians have both α- and βTRPMs, and saw independent expansion of βTRPM.

**Fig. S13.** Reconciled and rearranged maximum likelihood tree of TRPM sequences from Chordata (bold, excluding ray-finned fish) as in **Fig. S8**, with Xenacoelomorpha. Chordates have α- and βTRPMs, and both α- and βTRPMs expanded in vertebrates.

**Fig. S14.** Reconciled and rearranged maximum likelihood tree of TRPM sequences from Lophotrochozoa (bold) as in **Fig. S9**, with Xenacoelomorpha removed. Lophotrochozoans have both α- and βTRPMs, and saw independent expansion of βTRPM.

**Fig. S15.** Reconciled and rearranged maximum likelihood tree of TRPM sequences from Priapulida and Nematoda (bold) as in **Fig. S10**, with Xenacoelomorpha removed. Nematodes likely have both α- and βTRPMs, and likely saw expansion in both. However, given that *Toxocara canis* TRPMs are the only members of the nematode αTRPM clade, it is unclear if the α expansion was species-specific.

**Fig. S16.** Reconciled and rearranged maximum likelihood tree of TRPM sequences from Priapulida and Arthropoda (bold) as in **Fig. S11**, with Xenacoelomorpha removed. The majority of Arthropods only have αTRPM (many having only a single copy), but several chelicerates and crustaceans may have channels more distantly related to human, priapulid, and cnidarian βTRPMs.

**Fig. S17.** Vertebrate TRPM8 was independently lost in most vertebrate lineages, surviving only in the lobe-finned fish lineage (including tetrapods). Maximum likelihood tree of TRPM sequences from Chordata (including lancelets, tunicates, agnathans, sharks, coelecanth, tetrapods, and ray-finned fish), rooted in TRPS, with the 8 vertebrate TRPM clades labeled. Unlabeled clades are from invertebrate species. UFboot confidence is indicated by red-green color scale. Black indicates rearranged branches (<95 UFBoot).

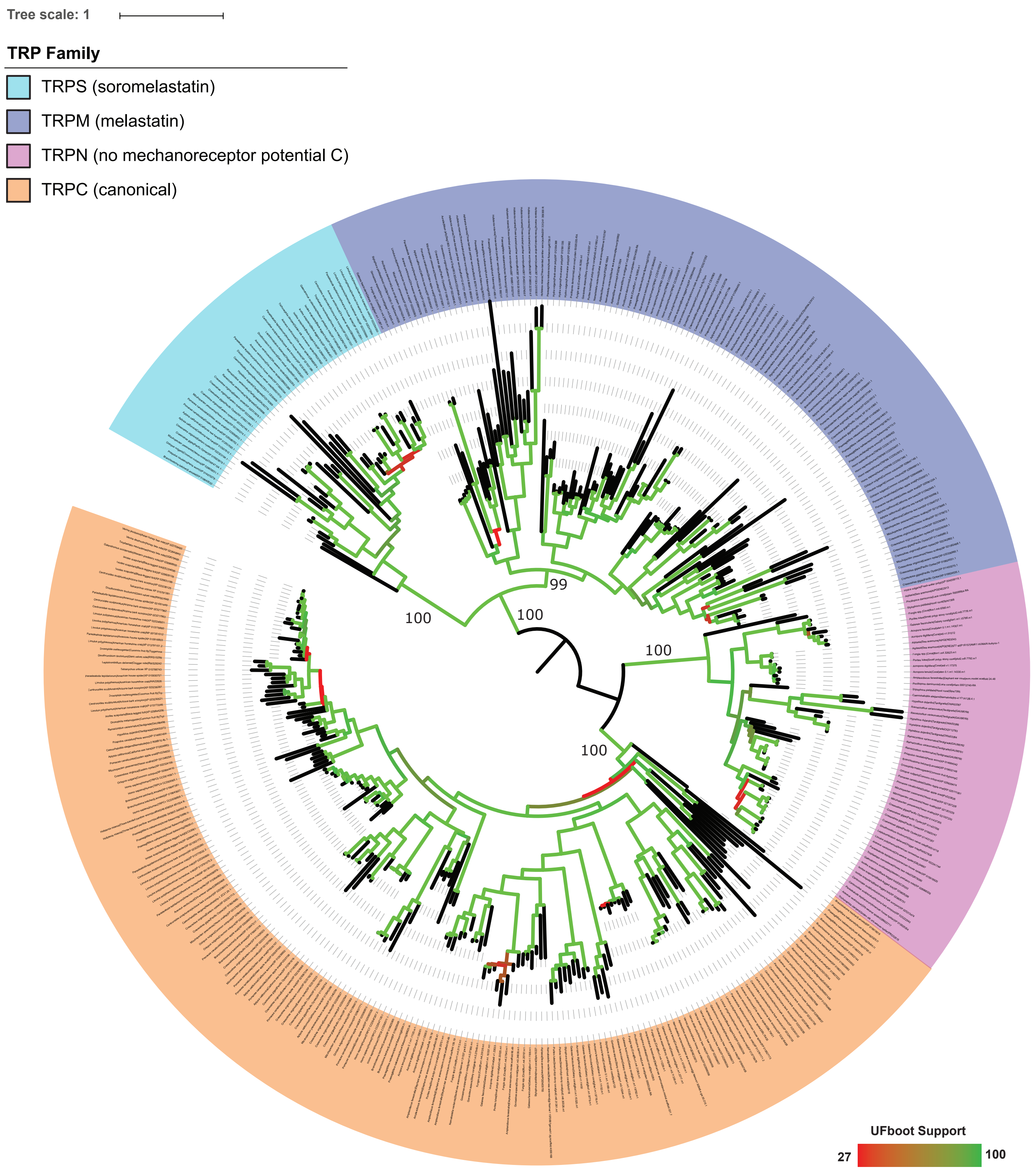
Tree scale: 1
TRP Family
TRPS (soromelastatin)
TRPM (melastatin)
TRPN (no mechanoreceptor potential C)
TRPC (canonical)
100
100
99
100
100
UFboot Support
27
100

Figure S1

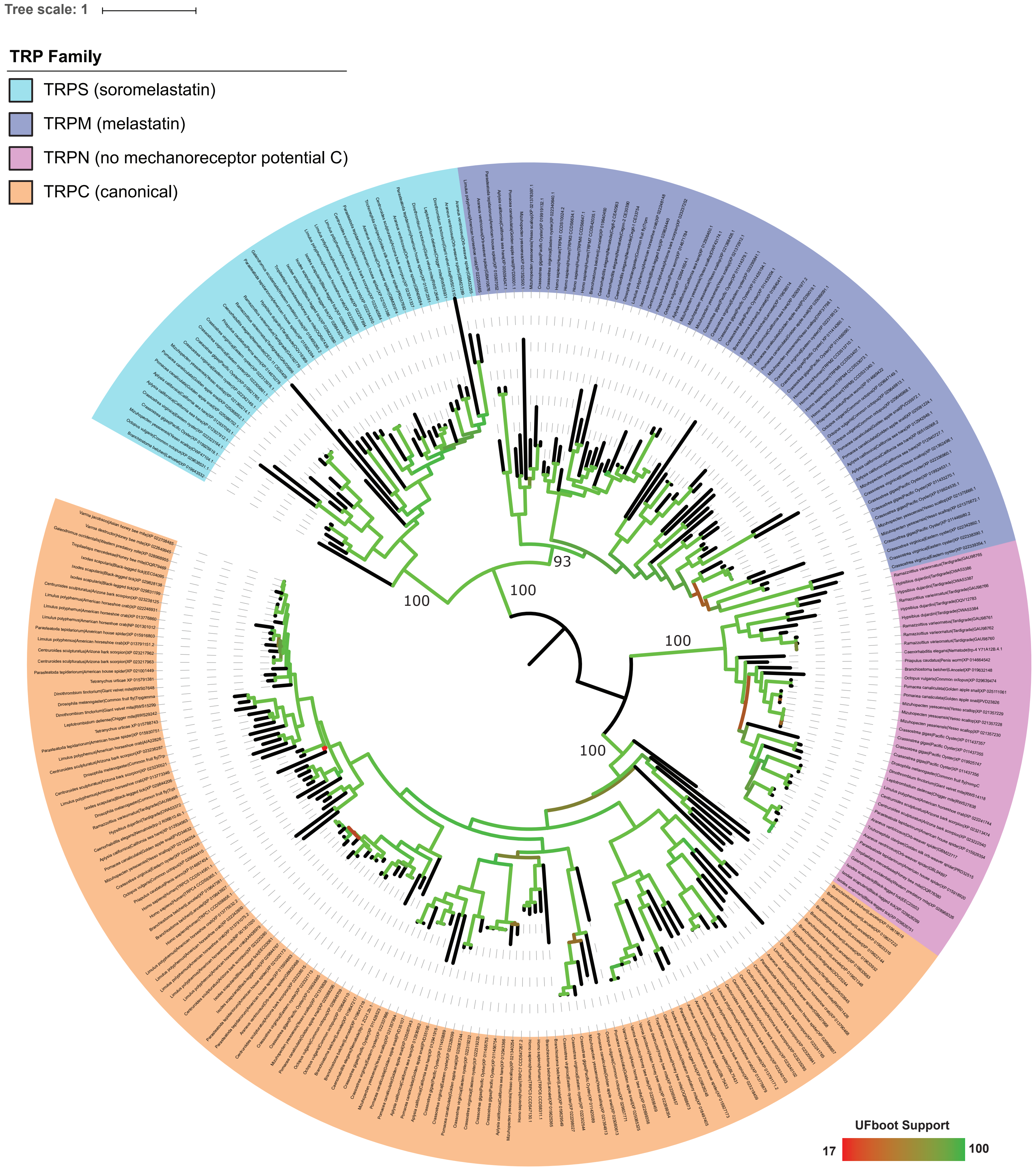
Tree scale: 1
TRP Family
TRPS (soromelastatin)
TRPM (melastatin)
TRPN (no mechanoreceptor potential C)
TRPC (canonical)
100
100
93
100
100
UFboot Support
17
100

Figure S2

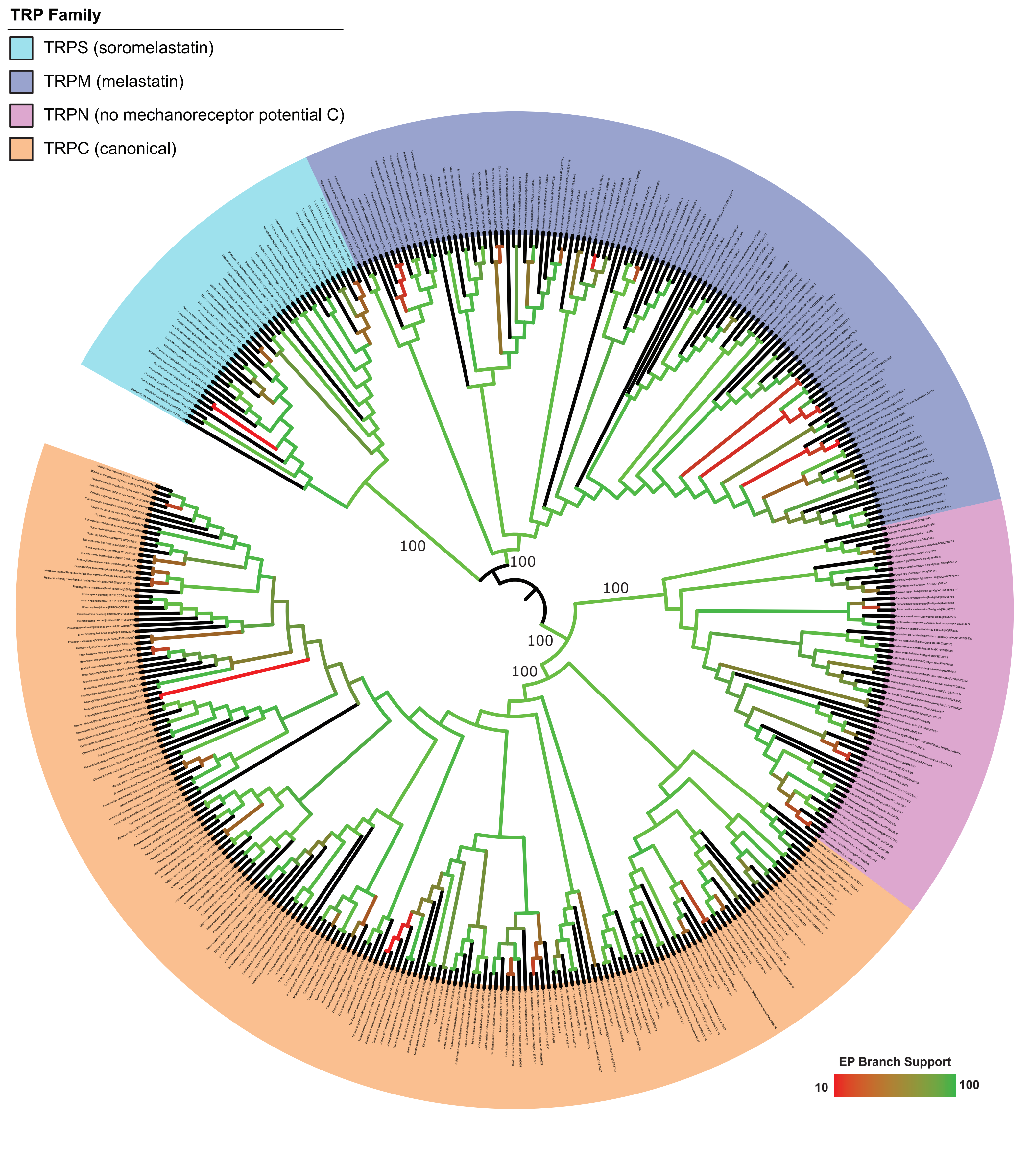

TRP Family
TRPS (soromelastatin)
TRPM (melastatin)
TRPN (no mechanoreceptor potential C)
TRPC (canonical)
100
100
100
100
100
EP Branch Support
10
100


Figure S3

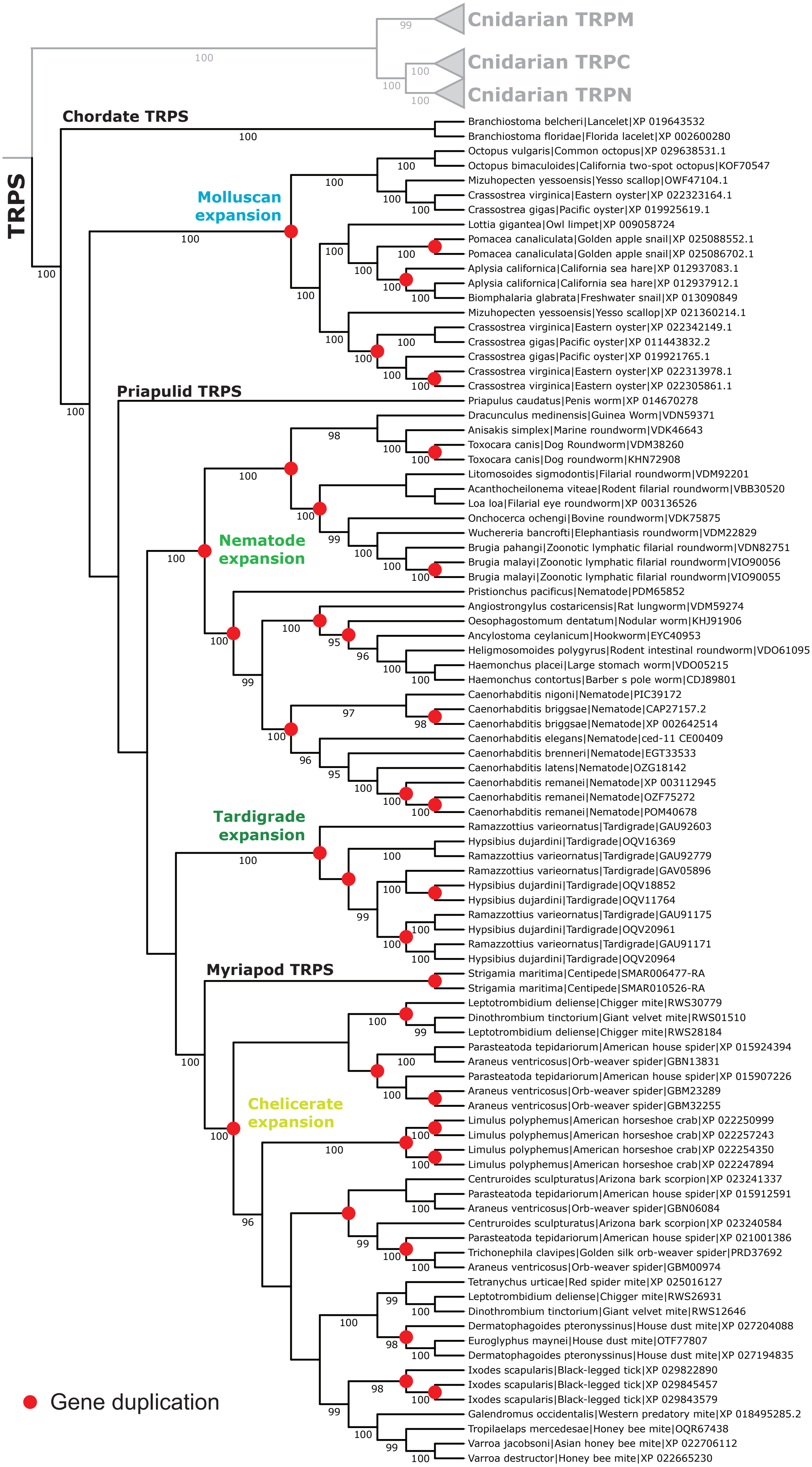
Cnidarian TRPM
Cnidarian TRPC
Cnidarian TRPN
TRPS
Chordate TRPS
Branchiostoma belcheri|Lancelet|XP 019643532
Branchiostoma floridae|Florida lacelet|XP 002600280
Molluscan expansion
Octopus vulgaris|Common octopus|XP 029638531.1
Octopus bimaculoides|California two-spot octopus|KOF70547
Mizuhopecten yessoensis|Yesso scallop|OWF47104.1
Crassostrea virginica|Eastern oyster|XP 022323164.1
Crassostrea gigas|Pacific oyster|XP 019925619.1
Lottia gigantea|Owl limpet|XP 009058724
Pomacea canaliculata|Golden apple snail|XP 025088552.1
Pomacea canaliculata|Golden apple snail|XP 025086702.1
Aplysia californica|California sea hare|XP 012937083.1
Aplysia californica|California sea hare|XP 012937912.1
Biomphalaria glabrata|Freshwater snail|XP 013090849
Mizuhopecten yessoensis|Yesso scallop|XP 021360214.1
Crassostrea virginica|Eastern oyster|XP 022342149.1
Crassostrea gigas|Pacific oyster|XP 011443832.2
Crassostrea gigas|Pacific oyster|XP 019921765.1
Crassostrea virginica|Eastern oyster|XP 022313978.1
Crassostrea virginica|Eastern oyster|XP 022305861.1
Priapulid TRPS
Priapulus caudatus|Penis worm|XP 014670278
Nematode expansion
Dracunculus medinensis|Guinea Worm|VDN59371
Anisakis simplex|Marine roundworm|VDK46643
Toxocara canis|Dog Roundworm|VDM38260
Toxocara canis|Dog roundworm|KHN72908
Litomosoides sigmodontis|Filarial roundworm|VDM92201
Acanthocheilonema viteae|Rodent filarial roundworm|VBB30520
Loa loa|Filarial eye roundworm|XP 003136526
Onchocerca ochengi|Bovine roundworm|VDK75875
Wuchereria bancrofti|Elephantiasis roundworm|VDM22829
Brugia pahangi|Zoonotic lymphatic filarial roundworm|VDN82751
Brugia malayi|Zoonotic lymphatic filarial roundworm|VIO90056
Brugia malayi|Zoonotic lymphatic filarial roundworm|VIO90055
Pristionchus pacificus|Nematode|PDM65852
Angiostrongylus costaricensis|Rat lungworm|VDM59274
Oesophagostomum dentatum|Nodular worm|KHJ91906
Ancylostoma ceylanicum|Hookworm|EYC40953
Heligmosomoides polygyrus|Rodent intestinal roundworm|VDO61095
Haemonchus placei|Large stomach worm|VDO05215
Haemonchus contortus|Barber s pole worm|CDJ89801
Caenorhabditis nigoni|Nematode|PIC39172
Caenorhabditis briggsae|Nematode|CAP27157.2
Caenorhabditis briggsae|Nematode|XP 002642514
Caenorhabditis elegans|Nematode|ced-11 CE00409
Caenorhabditis brenneri|Nematode|EGT33533
Caenorhabditis latens|Nematode|OZG18142
Caenorhabditis remanei|Nematode|XP 003112945
Caenorhabditis remanei|Nematode|OZF75272
Caenorhabditis remanei|Nematode|POM40678
Tardigrade expansion
Ramazzottius varieornatus|Tardigrade|GAU92603
Hypsibius dujardini|Tardigrade|OQV16369
Ramazzottius varieornatus|Tardigrade|GAU92779
Ramazzottius varieornatus|Tardigrade|GAV05896
Hypsibius dujardini|Tardigrade|OQV18852
Hypsibius dujardini|Tardigrade|OQV11764
Ramazzottius varieornatus|Tardigrade|GAU91175
Hypsibius dujardini|Tardigrade|OQV20961
Ramazzottius varieornatus|Tardigrade|GAU91171
Hypsibius dujardini|Tardigrade|OQV20964
Myriapod TRPS
Strigamia maritima|Centipede|SMAR006477-RA
Strigamia maritima|Centipede|SMAR010526-RA
Chelicerate expansion
Leptotrombidium deliense|Chigger mite|RWS30779
Dinothrombium tinctorium|Giant velvet mite|RWS01510
Leptotrombidium deliense|Chigger mite|RWS28184
Parasteatoda tepidariorum|American house spider|XP 015924394
Araneus ventricosus|Orb-weaver spider|GBN13831
Parasteatoda tepidariorum|American house spider|XP 015907226
Araneus ventricosus|Orb-weaver spider|GBM23289
Araneus ventricosus|Orb-weaver spider|GBM32255
Limulus polyphemus|American horseshoe crab|XP 022250999
Limulus polyphemus|American horseshoe crab|XP 022257243
Limulus polyphemus|American horseshoe crab|XP 022254350
Limulus polyphemus|American horseshoe crab|XP 022247894
Centruroides sculpturatus|Arizona bark scorpion|XP 023241337
Parasteatoda tepidariorum|American house spider|XP 015912591
Araneus ventricosus|Orb-weaver spider|GBN06084
Centruroides sculpturatus|Arizona bark scorpion|XP 023240584
Parasteatoda tepidariorum|American house spider|XP 021001386
Trichonephila clavipes|Golden silk orb-weaver spider|PRD37692
Araneus ventricosus|Orb-weaver spider|GBM00974
Tetranychus urticae|Red spider mite|XP 025016127
Leptotrombidium deliense|Chigger mite|RWS26931
Dinothrombium tinctorium|Giant velvet mite|RWS12646
Dermatophagoides pteronyssinus|House dust mite|XP 027204088
Euroglyphus maynei|House dust mite|OTF77807
Dermatophagoides pteronyssinus|House dust mite|XP 027194835
Ixodes scapularis|Black-legged tick|XP 029822890
Ixodes scapularis|Black-legged tick|XP 029845457
Ixodes scapularis|Black-legged tick|XP 029843579
Galendromus occidentalis|Western predatory mite|XP 018495285.2
Tropilaelaps mercedesae|Honey bee mite|OQR67438
Varroa jacobsoni|Asian honey bee mite|XP 022706112
Varroa destructor|Honey bee mite|XP 022665230
Gene duplication

Figure S4

#### Monophyletic Deuterostomia Hypothesis

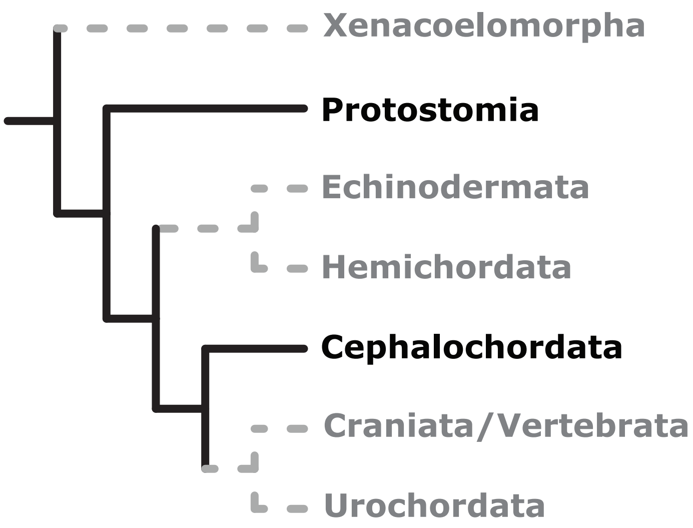


#### Xenambulacraria Hypothesis

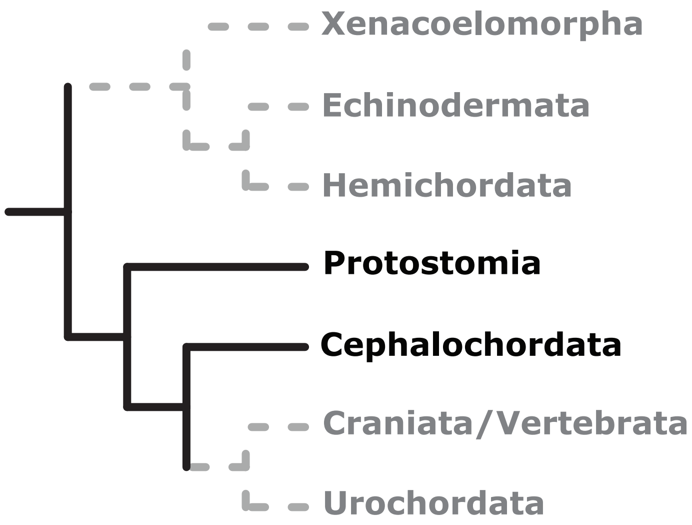


Figure S5

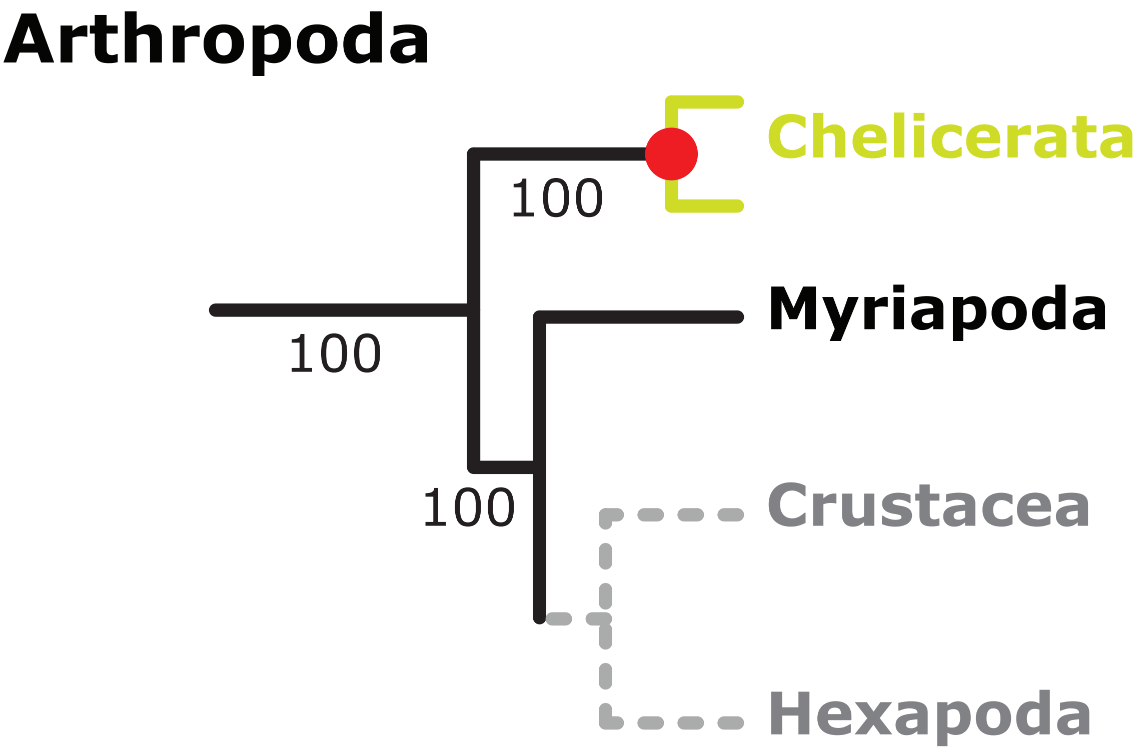
Arthropoda
Chelicerata
100
Myriapoda
100
100
Crustacea
Hexapoda

Figure S6

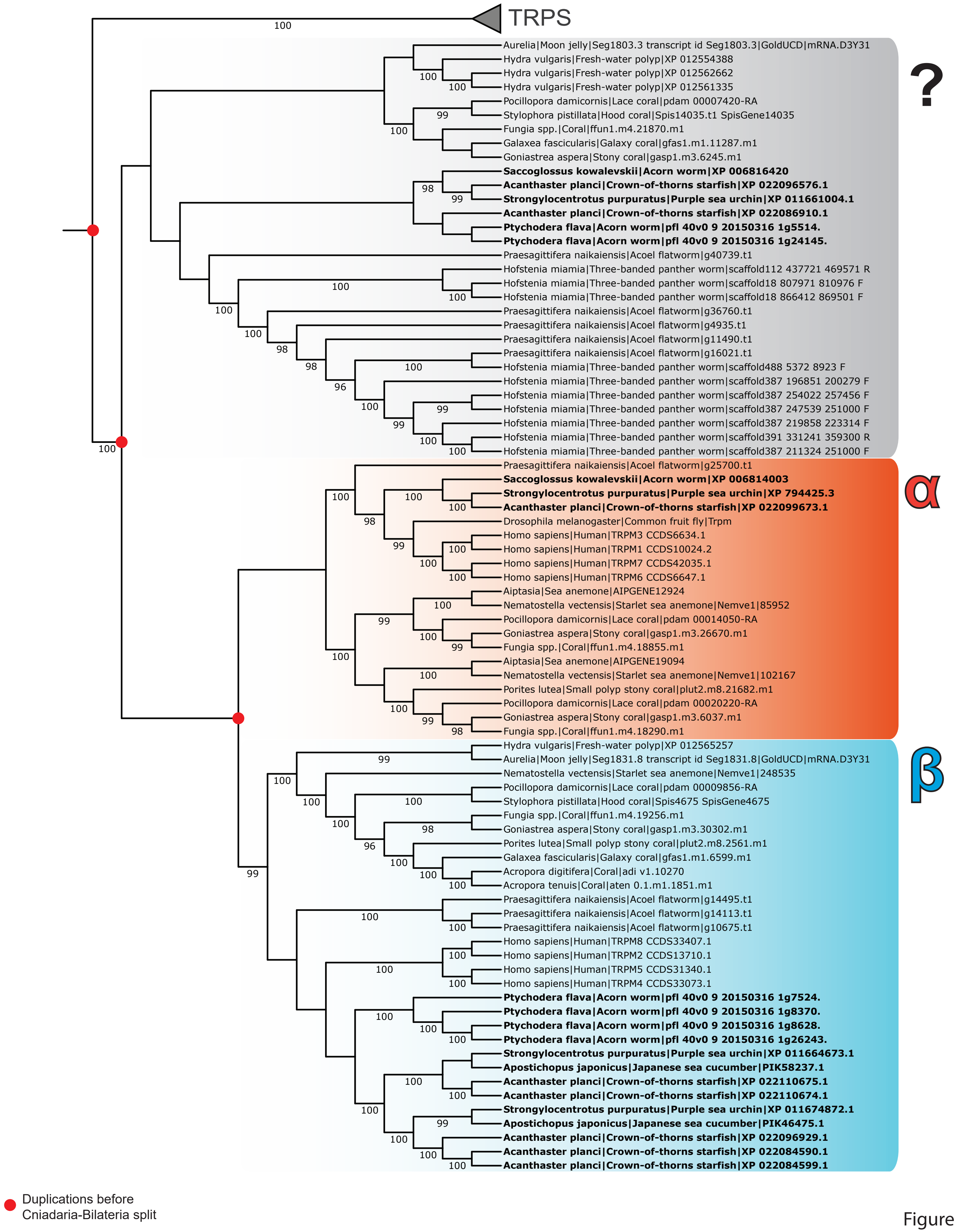
TRPS
?
Aurelia|Moon jelly|Seg1803.3 transcript id Seg1803.3|GoldUCD|mRNA.D3Y31
Hydra vulgaris|Fresh-water polyp|XP 012554388
Hydra vulgaris|Fresh-water polyp|XP 012562662
Hydra vulgaris|Fresh-water polyp|XP 012561335
Pocillopora damicornis|Lace coral|pdam 00007420-RA
Stylophora pistillata|Hood coral|Spis14035.t1 SpisGene14035
Fungia spp.|Coral|ffun1.m4.21870.m1
Galaxea fascicularis|Galaxy coral|gfas1.m1.11287.m1
Goniastrea aspera|Stony coral|gasp1.m3.6245.m1
Saccoglossus kowalevskii|Acorn worm|XP 006816420
Acanthaster planci|Crown-of-thorns starfish|XP 022096576.1
Strongylocentrotus purpuratus|Purple sea urchin|XP 011661004.1
Acanthaster planci|Crown-of-thorns starfish|XP 022086910.1
Ptychodera flava|Acorn worm|pfl 40v0 9 20150316 1g5514.
Ptychodera flava|Acorn worm|pfl 40v0 9 20150316 1g24145.
Praesagittifera naikaiensis|Acoel flatworm|g40739.t1
Hofstenia miamia|Three-banded panther worm|scaffold112 437721 469571 R
Hofstenia miamia|Three-banded panther worm|scaffold18 807971 810976 F
Hofstenia miamia|Three-banded panther worm|scaffold18 866412 869501 F
Praesagittifera naikaiensis|Acoel flatworm|g36760.t1
Praesagittifera naikaiensis|Acoel flatworm|g4935.t1
Praesagittifera naikaiensis|Acoel flatworm|g11490.t1
Praesagittifera naikaiensis|Acoel flatworm|g16021.t1
Hofstenia miamia|Three-banded panther worm|scaffold488 5372 8923 F
Hofstenia miamia|Three-banded panther worm|scaffold387 196851 200279 F
Hofstenia miamia|Three-banded panther worm|scaffold387 254022 257456 F
Hofstenia miamia|Three-banded panther worm|scaffold387 247539 251000 F
Hofstenia miamia|Three-banded panther worm|scaffold387 219858 223314 F
Hofstenia miamia|Three-banded panther worm|scaffold391 331241 359300 R
Hofstenia miamia|Three-banded panther worm|scaffold387 211324 251000 F
α
Praesagittifera naikaiensis|Acoel flatworm|g25700.t1
Saccoglossus kowalevskii|Acorn worm|XP 006814003
Strongylocentrotus purpuratus|Purple sea urchin|XP 794425.3
Acanthaster planci|Crown-of-thorns starfish|XP 022099673.1
Drosophila melanogaster|Common fruit fly|Trpm
Homo sapiens|Human|TRPM3 CCDS6634.1
Homo sapiens|Human|TRPM1 CCDS10024.2
Homo sapiens|Human|TRPM7 CCDS42035.1
Homo sapiens|Human|TRPM6 CCDS6647.1
Aiptasia|Sea anemone|AIPGENE12924
Nematostella vectensis|Starlet sea anemone|Nemve1|85952
Pocillopora damicornis|Lace coral|pdam 00014050-RA
Goniastrea aspera|Stony coral|gasp1.m3.26670.m1
Fungia spp.|Coral|ffun1.m4.18855.m1
Aiptasia|Sea anemone|AIPGENE19094
Nematostella vectensis|Starlet sea anemone|Nemve1|102167
Porites lutea|Small polyp stony coral|plut2.m8.21682.m1
Pocillopora damicornis|Lace coral|pdam 00020220-RA
Goniastrea aspera|Stony coral|gasp1.m3.6037.m1
Fungia spp.|Coral|ffun1.m4.18290.m1
β
Hydra vulgaris|Fresh-water polyp|XP 012565257
Aurelia|Moon jelly|Seg1831.8 transcript id Seg1831.8|GoldUCD|mRNA.D3Y31
Nematostella vectensis|Starlet sea anemone|Nemve1|248535
Pocillopora damicornis|Lace coral|pdam 00009856-RA
Stylophora pistillata|Hood coral|Spis4675 SpisGene4675
Fungia spp.|Coral|ffun1.m4.19256.m1
Goniastrea aspera|Stony coral|gasp1.m3.30302.m1
Porites lutea|Small polyp stony coral|plut2.m8.2561.m1
Galaxea fascicularis|Galaxy coral|gfas1.m1.6599.m1
Acropora digitifera|Coral|adi v1.10270
Acropora tenuis|Coral|aten 0.1.m1.1851.m1
Praesagittifera naikaiensis|Acoel flatworm|g14495.t1
Praesagittifera naikaiensis|Acoel flatworm|g14113.t1
Praesagittifera naikaiensis|Acoel flatworm|g10675.t1
Homo sapiens|Human|TRPM8 CCDS33407.1
Homo sapiens|Human|TRPM2 CCDS13710.1
Homo sapiens|Human|TRPM5 CCDS31340.1
Homo sapiens|Human|TRPM4 CCDS33073.1
Ptychodera flava|Acorn worm|pfl 40v0 9 20150316 1g7524.
Ptychodera flava|Acorn worm|pfl 40v0 9 20150316 1g8370.
Ptychodera flava|Acorn worm|pfl 40v0 9 20150316 1g8628.
Ptychodera flava|Acorn worm|pfl 40v0 9 20150316 1g26243.
Strongylocentrotus purpuratus|Purple sea urchin|XP 011664673.1
Apostichopus japonicus|Japanese sea cucumber|PIK58237.1
Acanthaster planci|Crown-of-thorns starfish|XP 022110675.1
Acanthaster planci|Crown-of-thorns starfish|XP 022110674.1
Strongylocentrotus purpuratus|Purple sea urchin|XP 011674872.1
Apostichopus japonicus|Japanese sea cucumber|PIK46475.1
Acanthaster planci|Crown-of-thorns starfish|XP 022096929.1
Acanthaster planci|Crown-of-thorns starfish|XP 022084590.1
Acanthaster planci|Crown-of-thorns starfish|XP 022084599.1
Duplications before Cniadaria-Bilateria split

Figure S7

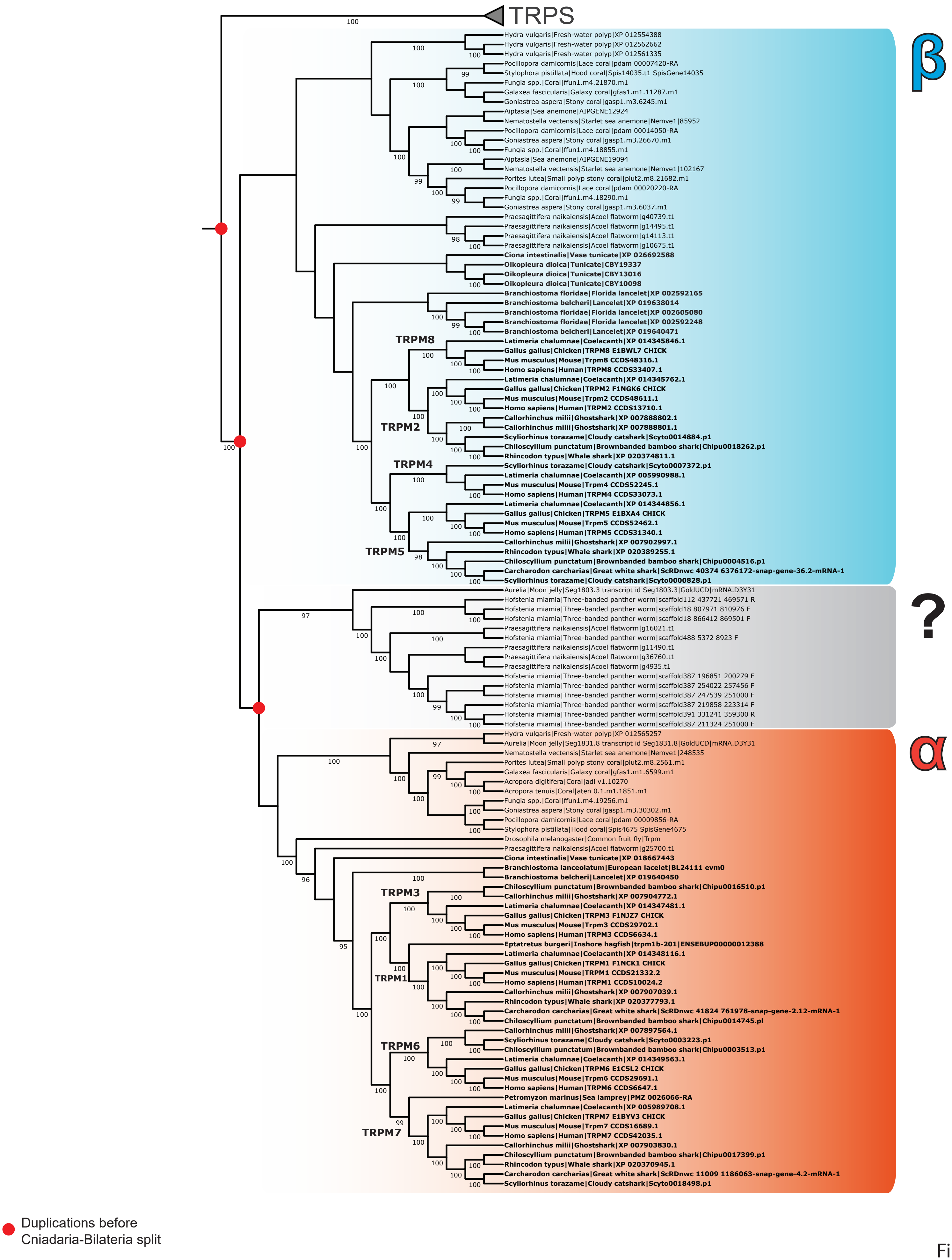

TRPS
β
Hydra vulgaris|Fresh-water polyp|XP 012554388
Hydra vulgaris|Fresh-water polyp|XP 012562662
Hydra vulgaris|Fresh-water polyp|XP 012561335
Pocillopora damicornis|Lace coral|pdam 00007420-RA
Stylophora pistillata|Hood coral|Spis14035.t1 SpisGene14035
Fungia spp.|Coral|ffun1.m4.21870.m1
Galaxea fascicularis|Galaxy coral|gfas1.m1.11287.m1
Goniastrea aspera|Stony coral|gasp1.m3.6245.m1
Aiptasia|Sea anemone|AIPGENE12924
Nematostella vectensis|Starlet sea anemone|Nemve1|85952
Pocillopora damicornis|Lace coral|pdam 00014050-RA
Goniastrea aspera|Stony coral|gasp1.m3.26670.m1
Fungia spp.|Coral|ffun1.m4.18855.m1
Aiptasia|Sea anemone|AIPGENE19094
Nematostella vectensis|Starlet sea anemone|Nemve1|102167
Porites lutea|Small polyp stony coral|plut2.m8.21682.m1
Pocillopora damicornis|Lace coral|pdam 00020220-RA
Fungia spp.|Coral|ffun1.m4.18290.m1
Goniastrea aspera|Stony coral|gasp1.m3.6037.m1
Praesagittifera naikaiensis|Acoel flatworm|g40739.t1
Praesagittifera naikaiensis|Acoel flatworm|g14495.t1
Praesagittifera naikaiensis|Acoel flatworm|g14113.t1
Praesagittifera naikaiensis|Acoel flatworm|g10675.t1
Ciona intestinalis|Vase tunicate|XP 026692588
Oikopleura dioica|Tunicate|CBY19337
Oikopleura dioica|Tunicate|CBY13016
Oikopleura dioica|Tunicate|CBY10098
Branchiostoma floridae|Florida lancelet|XP 002592165
Branchiostoma belcheri|Lancelet|XP 019638014
Branchiostoma floridae|Florida lancelet|XP 002605080
Branchiostoma floridae|Florida lancelet|XP 002592248
Branchiostoma belcheri|Lancelet|XP 019640471
TRPM8
Latimeria chalumnae|Coelacanth|XP 014345846.1
Gallus gallus|Chicken|TRPM8 E1BWL7 CHICK
Mus musculus|Mouse|Trpm8 CCDS48316.1
Homo sapiens|Human|TRPM8 CCDS33407.1
Latimeria chalumnae|Coelacanth|XP 014345762.1
Gallus gallus|Chicken|TRPM2 F1NGK6 CHICK
Mus musculus|Mouse|Trpm2 CCDS48611.1
Homo sapiens|Human|TRPM2 CCDS13710.1
TRPM2
Callorhinchus milii|Ghostshark|XP 007888802.1
Callorhinchus milii|Ghostshark|XP 007888801.1
Scyliorhinus torazame|Cloudy catshark|Scyto0014884.p1
Chiloscyllium punctatum|Brownbanded bamboo shark|Chipu0018262.p1
Rhincodon typus|Whale shark|XP 020374811.1
TRPM4
Scyliorhinus torazame|Cloudy catshark|Scyto0007372.p1
Latimeria chalumnae|Coelacanth|XP 005990988.1
Mus musculus|Mouse|Trpm4 CCDS52245.1
Homo sapiens|Human|TRPM4 CCDS33073.1
Latimeria chalumnae|Coelacanth|XP 014344856.1
Gallus gallus|Chicken|TRPM5 E1BXA4 CHICK
Mus musculus|Mouse|Trpm5 CCDS52462.1
Homo sapiens|Human|TRPM5 CCDS31340.1
Callorhinchus milii|Ghostshark|XP 007902997.1
TRPM5
Rhincodon typus|Whale shark|XP 020389255.1
Chiloscyllium punctatum|Brownbanded bamboo shark|Chipu0004516.p1
Carcharodon carcharias|Great white shark|ScRDnwc 40374 6376172-snap-gene-36.2-mRNA-1
Scyliorhinus torazame|Cloudy catshark|Scyto0000828.p1
?
Aurelia|Moon jelly|Seg1803.3 transcript id Seg1803.3|GoldUCD|mRNA.D3Y31
Hofstenia miamia|Three-banded panther worm|scaffold112 437721 469571 R
Hofstenia miamia|Three-banded panther worm|scaffold18 807971 810976 F
Hofstenia miamia|Three-banded panther worm|scaffold18 866412 869501 F
Praesagittifera naikaiensis|Acoel flatworm|g16021.t1
Hofstenia miamia|Three-banded panther worm|scaffold488 5372 8923 F
Praesagittifera naikaiensis|Acoel flatworm|g11490.t1
Praesagittifera naikaiensis|Acoel flatworm|g36760.t1
Praesagittifera naikaiensis|Acoel flatworm|g4935.t1
Hofstenia miamia|Three-banded panther worm|scaffold387 196851 200279 F
Hofstenia miamia|Three-banded panther worm|scaffold387 254022 257456 F
Hofstenia miamia|Three-banded panther worm|scaffold387 247539 251000 F
Hofstenia miamia|Three-banded panther worm|scaffold387 219858 223314 F
Hofstenia miamia|Three-banded panther worm|scaffold391 331241 359300 R
Hofstenia miamia|Three-banded panther worm|scaffold387 211324 251000 F
α
Hydra vulgaris|Fresh-water polyp|XP 012565257
Aurelia|Moon jelly|Seg1831.8 transcript id Seg1831.8|GoldUCD|mRNA.D3Y31
Nematostella vectensis|Starlet sea anemone|Nemve1|248535
Porites lutea|Small polyp stony coral|plut2.m8.2561.m1
Galaxea fascicularis|Galaxy coral|gfas1.m1.6599.m1
Acropora digitifera|Coral|adi v1.10270
Acropora tenuis|Coral|aten 0.1.m1.1851.m1
Fungia spp.|Coral|ffun1.m4.19256.m1
Goniastrea aspera|Stony coral|gasp1.m3.30302.m1
Pocillopora damicornis|Lace coral|pdam 00009856-RA
Stylophora pistillata|Hood coral|Spis4675 SpisGene4675
Drosophila melanogaster|Common fruit fly|Trpm
Praesagittifera naikaiensis|Acoel flatworm|g25700.t1
Ciona intestinalis|Vase tunicate|XP 018667443
Branchiostoma lanceolatum|European lacelet|BL24111 evm0
Branchiostoma belcheri|Lancelet|XP 019640450
TRPM3
Chiloscyllium punctatum|Brownbanded bamboo shark|Chipu0016510.p1
Callorhinchus milii|Ghostshark|XP 007904772.1
Latimeria chalumnae|Coelacanth|XP 014347481.1
Gallus gallus|Chicken|TRPM3 F1NJZ7 CHICK
Mus musculus|Mouse|Trpm3 CCDS29702.1
Homo sapiens|Human|TRPM3 CCDS6634.1
Eptatretus burgeri|Inshore hagfish|trpm1b-201|ENSEBUP00000012388
Latimeria chalumnae|Coelacanth|XP 014348116.1
Gallus gallus|Chicken|TRPM1 F1NCK1 CHICK
Mus musculus|Mouse|TRPM1 CCDS21332.2
TRPM1
Homo sapiens|Human|TRPM1 CCDS10024.2
Callorhinchus milii|Ghostshark|XP 007907039.1
Rhincodon typus|Whale shark|XP 020377793.1
Carcharodon carcharias|Great white shark|ScRDnwc 41824 761978-snap-gene-2.12-mRNA-1
Chiloscyllium punctatum|Brownbanded bamboo shark|Chipu0014745.pl
Callorhinchus milii|Ghostshark|XP 007897564.1
Scyliorhinus torazame|Cloudy catshark|Scyto0003223.p1
TRPM6
Chiloscyllium punctatum|Brownbanded bamboo shark|Chipu0003513.p1
Latimeria chalumnae|Coelacanth|XP 014349563.1
Gallus gallus|Chicken|TRPM6 E1C5L2 CHICK
Mus musculus|Mouse|Trpm6 CCDS29691.1
Homo sapiens|Human|TRPM6 CCDS6647.1
Petromyzon marinus|Sea lamprey|PMZ 0026066-RA
Latimeria chalumnae|Coelacanth|XP 005989708.1
Gallus gallus|Chicken|TRPM7 E1BYV3 CHICK
Mus musculus|Mouse|Trpm7 CCDS16689.1
TRPM7
Homo sapiens|Human|TRPM7 CCDS42035.1
Callorhinchus milii|Ghostshark|XP 007903830.1
Chiloscyllium punctatum|Brownbanded bamboo shark|Chipu0017399.p1
Rhincodon typus|Whale shark|XP 020370945.1
Carcharodon carcharias|Great white shark|ScRDnwc 11009 1186063-snap-gene-4.2-mRNA-1
Scyliorhinus torazame|Cloudy catshark|Scyto0018498.p1
Duplications before
Cniadaria-Bilateria split


Figure S8

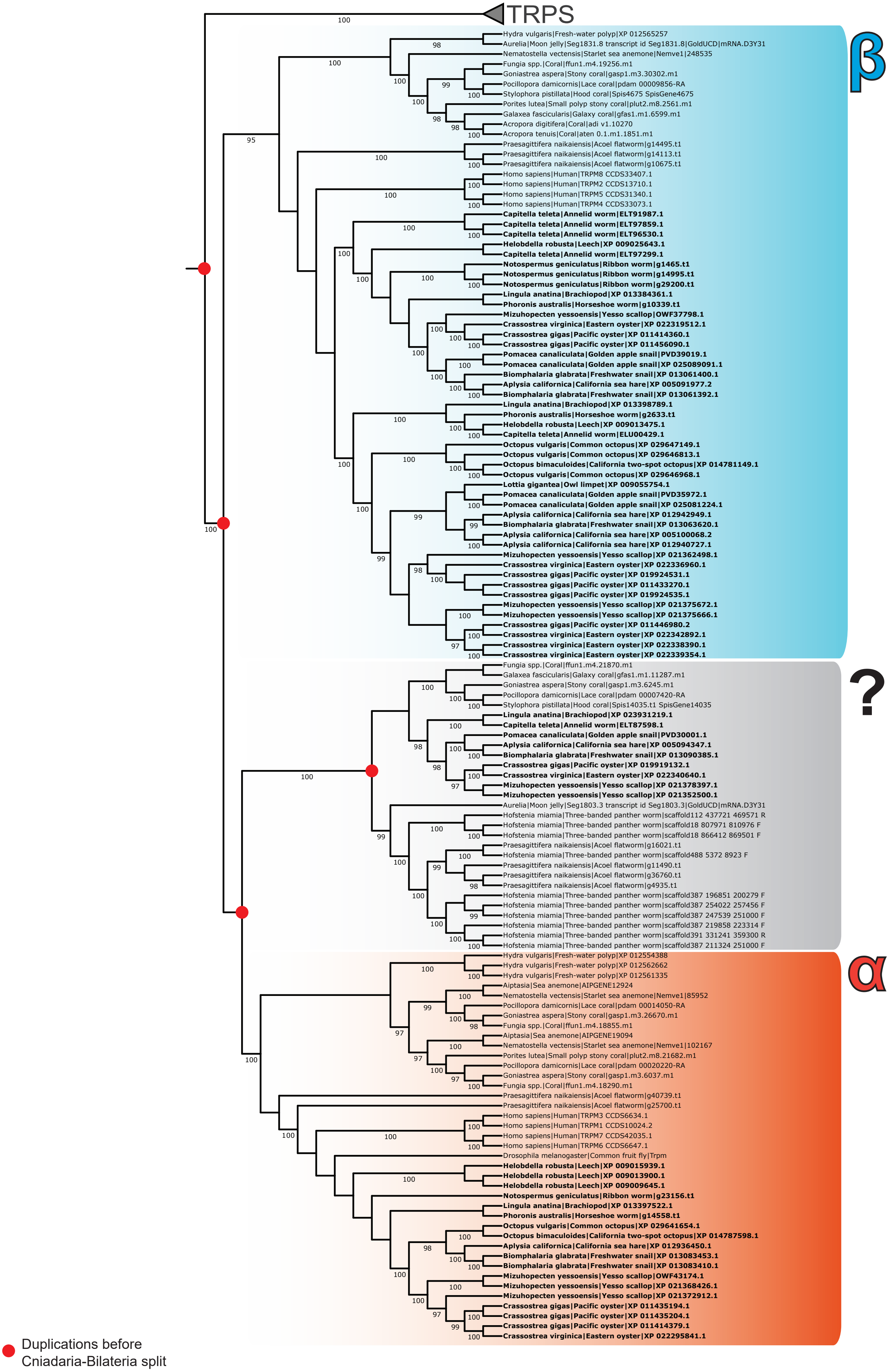

TRPS
β
Hydra vulgaris|Fresh-water polyp|XP 012565257
Aurelia|Moon jelly|Seg1831.8 transcript id Seg1831.8|GoldUCD|mRNA.D3Y31
Nematostella vectensis|Starlet sea anemone|Nemve1|248535
Fungia spp.|Coral|ffun1.m4.19256.m1
Goniastrea aspera|Stony coral|gasp1.m3.30302.m1
Pocillopora damicornis|Lace coral|pdam 00009856-RA
Stylophora pistillata|Hood coral|Spis4675 SpisGene4675
Porites lutea|Small polyp stony coral|plut2.m8.2561.m1
Galaxea fascicularis|Galaxy coral|gfas1.m1.6599.m1
Acropora digitifera|Coral|adi v1.10270
Acropora tenuis|Coral|aten 0.1.m1.1851.m1
Praesagittifera naikaiensis|Acoel flatworm|g14495.t1
Praesagittifera naikaiensis|Acoel flatworm|g14113.t1
Praesagittifera naikaiensis|Acoel flatworm|g10675.t1
Homo sapiens|Human|TRPM8 CCDS33407.1
Homo sapiens|Human|TRPM2 CCDS13710.1
Homo sapiens|Human|TRPM5 CCDS31340.1
Homo sapiens|Human|TRPM4 CCDS33073.1
Capitella teleta|Annelid worm|ELT91987.1
Capitella teleta|Annelid worm|ELT97859.1
Capitella teleta|Annelid worm|ELT96530.1
Helobdella robusta|Leech|XP 009025643.1
Capitella teleta|Annelid worm|ELT97299.1
Notospermus geniculatus|Ribbon worm|g1465.t1
Notospermus geniculatus|Ribbon worm|g14995.t1
Notospermus geniculatus|Ribbon worm|g29200.t1
Lingula anatina|Brachiopod|XP 013384361.1
Phoronis australis|Horseshoe worm|g10339.t1
Mizuhopecten yessoensis|Yesso scallop|OWF37798.1
Crassostrea virginica|Eastern oyster|XP 022319512.1
Crassostrea gigas|Pacific oyster|XP 011414360.1
Crassostrea gigas|Pacific oyster|XP 011456090.1
Pomacea canaliculata|Golden apple snail|PVD39019.1
Pomacea canaliculata|Golden apple snail|XP 025089091.1
Biomphalaria glabrata|Freshwater snail|XP 013061400.1
Aplysia californica|California sea hare|XP 005091977.2
Biomphalaria glabrata|Freshwater snail|XP 013061392.1
Lingula anatina|Brachiopod|XP 013398789.1
Phoronis australis|Horseshoe worm|g2633.t1
Helobdella robusta|Leech|XP 009013475.1
Capitella teleta|Annelid worm|ELU00429.1
Octopus vulgaris|Common octopus|XP 029647149.1
Octopus vulgaris|Common octopus|XP 029646813.1
Octopus bimaculoides|California two-spot octopus|XP 014781149.1
Octopus vulgaris|Common octopus|XP 029646968.1
Lottia gigantea|Owl limpet|XP 009055754.1
Pomacea canaliculata|Golden apple snail|PVD35972.1
Pomacea canaliculata|Golden apple snail|XP 025081224.1
Aplysia californica|California sea hare|XP 012942949.1
Biomphalaria glabrata|Freshwater snail|XP 013063620.1
Aplysia californica|California sea hare|XP 005100068.2
Aplysia californica|California sea hare|XP 012940727.1
Mizuhopecten yessoensis|Yesso scallop|XP 021362498.1
Crassostrea virginica|Eastern oyster|XP 022336960.1
Crassostrea gigas|Pacific oyster|XP 019924531.1
Crassostrea gigas|Pacific oyster|XP 011433270.1
Crassostrea gigas|Pacific oyster|XP 019924535.1
Mizuhopecten yessoensis|Yesso scallop|XP 021375672.1
Mizuhopecten yessoensis|Yesso scallop|XP 021375666.1
Crassostrea gigas|Pacific oyster|XP 011446980.2
Crassostrea virginica|Eastern oyster|XP 022342892.1
Crassostrea virginica|Eastern oyster|XP 022338390.1
Crassostrea virginica|Eastern oyster|XP 022339354.1
?
Fungia spp.|Coral|ffun1.m4.21870.m1
Galaxea fascicularis|Galaxy coral|gfas1.m1.11287.m1
Goniastrea aspera|Stony coral|gasp1.m3.6245.m1
Pocillopora damicornis|Lace coral|pdam 00007420-RA
Stylophora pistillata|Hood coral|Spis14035.t1 SpisGene14035
Lingula anatina|Brachiopod|XP 023931219.1
Capitella teleta|Annelid worm|ELT87598.1
Pomacea canaliculata|Golden apple snail|PVD30001.1
Aplysia californica|California sea hare|XP 005094347.1
Biomphalaria glabrata|Freshwater snail|XP 013090385.1
Crassostrea gigas|Pacific oyster|XP 019919132.1
Crassostrea virginica|Eastern oyster|XP 022340640.1
Mizuhopecten yessoensis|Yesso scallop|XP 021378397.1
Mizuhopecten yessoensis|Yesso scallop|XP 021352500.1
Aurelia|Moon jelly|Seg1803.3 transcript id Seg1803.3|GoldUCD|mRNA.D3Y31
Hofstenia miamia|Three-banded panther worm|scaffold112 437721 469571 R
Hofstenia miamia|Three-banded panther worm|scaffold18 807971 810976 F
Hofstenia miamia|Three-banded panther worm|scaffold18 866412 869501 F
Praesagittifera naikaiensis|Acoel flatworm|g16021.t1
Hofstenia miamia|Three-banded panther worm|scaffold488 5372 8923 F
Praesagittifera naikaiensis|Acoel flatworm|g11490.t1
Praesagittifera naikaiensis|Acoel flatworm|g36760.t1
Praesagittifera naikaiensis|Acoel flatworm|g4935.t1
Hofstenia miamia|Three-banded panther worm|scaffold387 196851 200279 F
Hofstenia miamia|Three-banded panther worm|scaffold387 254022 257456 F
Hofstenia miamia|Three-banded panther worm|scaffold387 247539 251000 F
Hofstenia miamia|Three-banded panther worm|scaffold387 219858 223314 F
Hofstenia miamia|Three-banded panther worm|scaffold391 331241 359300 R
Hofstenia miamia|Three-banded panther worm|scaffold387 211324 251000 F
α
Hydra vulgaris|Fresh-water polyp|XP 012554388
Hydra vulgaris|Fresh-water polyp|XP 012562662
Hydra vulgaris|Fresh-water polyp|XP 012561335
Aiptasia|Sea anemone|AIPGENE12924
Nematostella vectensis|Starlet sea anemone|Nemve1|85952
Pocillopora damicornis|Lace coral|pdam 00014050-RA
Goniastrea aspera|Stony coral|gasp1.m3.26670.m1
Fungia spp.|Coral|ffun1.m4.18855.m1
Aiptasia|Sea anemone|AIPGENE19094
Nematostella vectensis|Starlet sea anemone|Nemve1|102167
Porites lutea|Small polyp stony coral|plut2.m8.21682.m1
Pocillopora damicornis|Lace coral|pdam 00020220-RA
Goniastrea aspera|Stony coral|gasp1.m3.6037.m1
Fungia spp.|Coral|ffun1.m4.18290.m1
Praesagittifera naikaiensis|Acoel flatworm|g40739.t1
Praesagittifera naikaiensis|Acoel flatworm|g25700.t1
Homo sapiens|Human|TRPM3 CCDS6634.1
Homo sapiens|Human|TRPM1 CCDS10024.2
Homo sapiens|Human|TRPM7 CCDS42035.1
Homo sapiens|Human|TRPM6 CCDS6647.1
Drosophila melanogaster|Common fruit fly|Trpm
Helobdella robusta|Leech|XP 009015939.1
Helobdella robusta|Leech|XP 009013900.1
Helobdella robusta|Leech|XP 009009645.1
Notospermus geniculatus|Ribbon worm|g23156.t1
Lingula anatina|Brachiopod|XP 013397522.1
Phoronis australis|Horseshoe worm|g14558.t1
Octopus vulgaris|Common octopus|XP 029641654.1
Octopus bimaculoides|California two-spot octopus|XP 014787598.1
Aplysia californica|California sea hare|XP 012936450.1
Biomphalaria glabrata|Freshwater snail|XP 013083453.1
Biomphalaria glabrata|Freshwater snail|XP 013083410.1
Mizuhopecten yessoensis|Yesso scallop|OWF43174.1
Mizuhopecten yessoensis|Yesso scallop|XP 021368426.1
Mizuhopecten yessoensis|Yesso scallop|XP 021372912.1
Crassostrea gigas|Pacific oyster|XP 011435194.1
Crassostrea gigas|Pacific oyster|XP 011435204.1
Crassostrea gigas|Pacific oyster|XP 011414379.1
Crassostrea virginica|Eastern oyster|XP 022295841.1
Duplications before
Cniadaria-Bilateria split


Figure S9

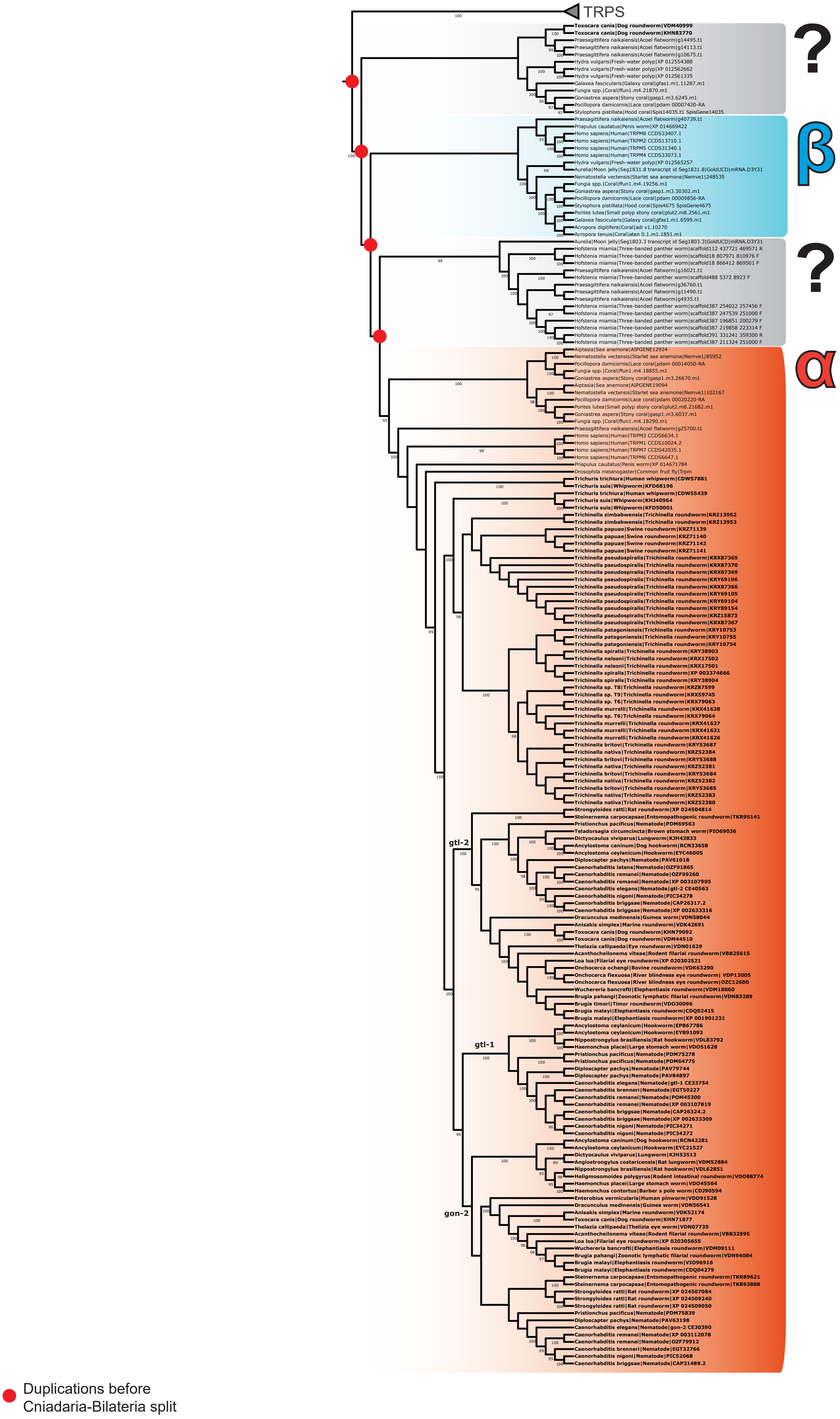

TRPS
?
β
?
α
gtl-2
gtl-1
gon-2
Duplications before
Cniadaria-Bilateria split


Figure S10

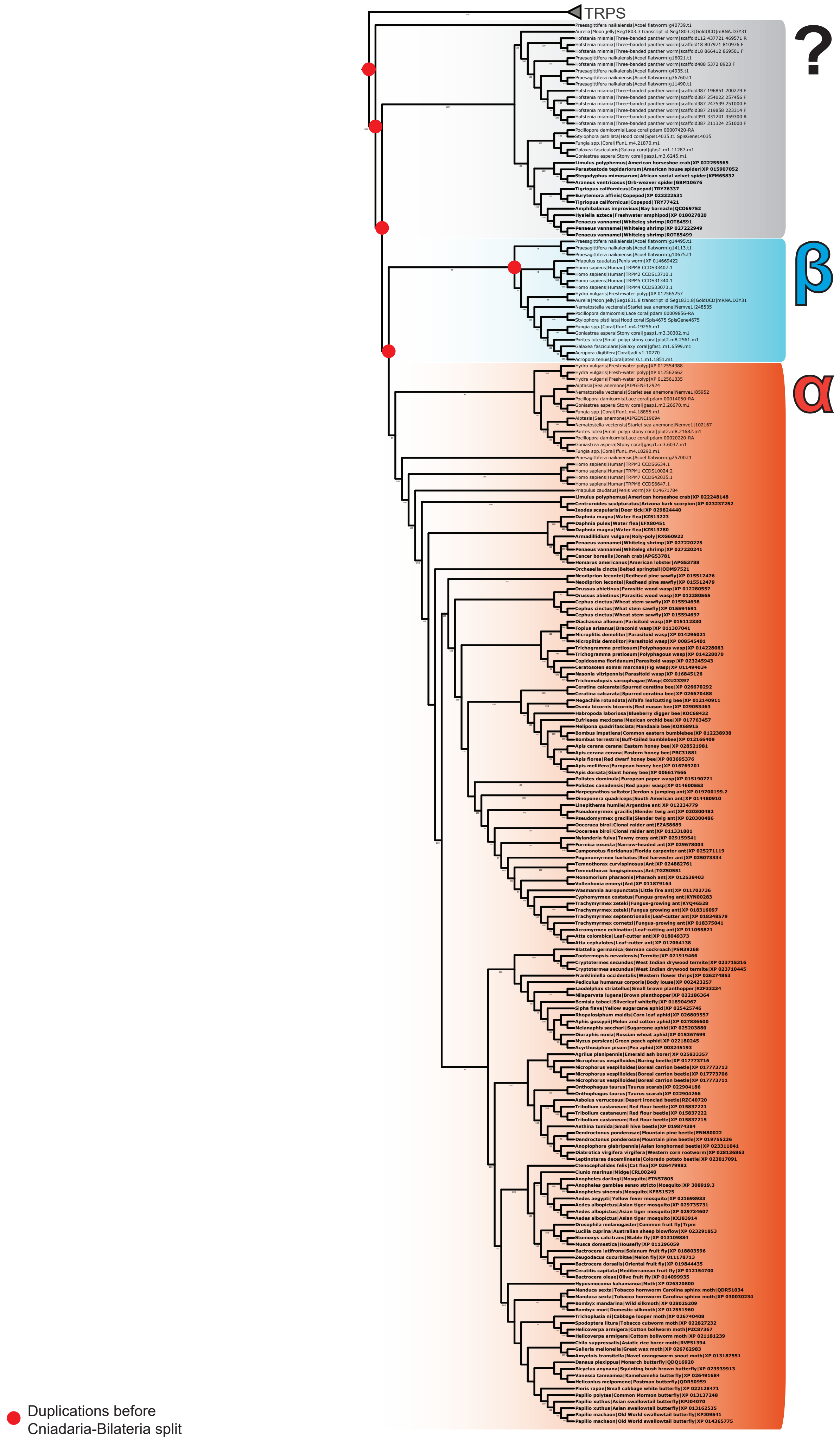


Figure S11

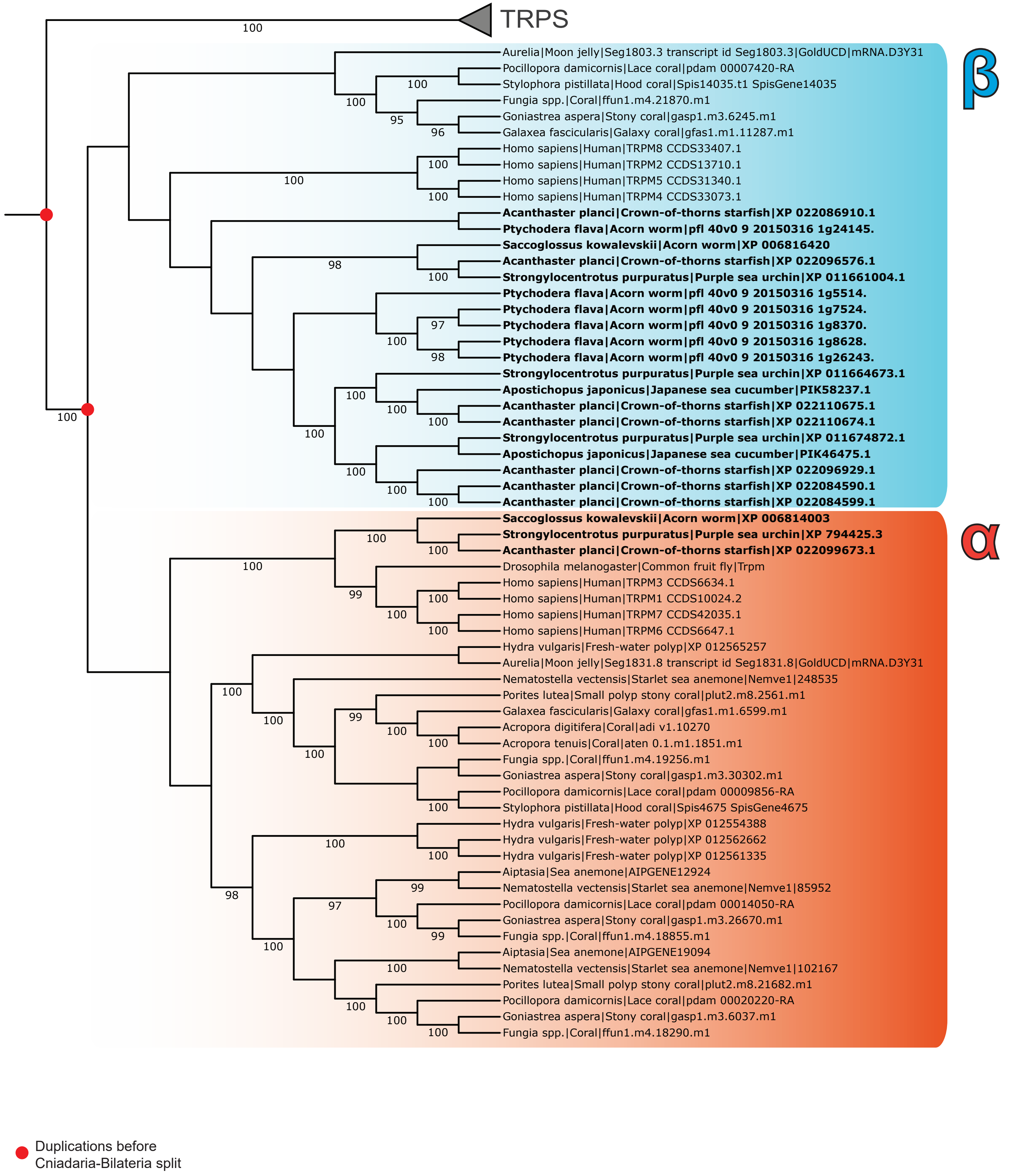
TRPS
β
Aurelia|Moon jelly|Seg1803.3 transcript id Seg1803.3|GoldUCD|mRNA.D3Y31
Pocillopora damicornis|Lace coral|pdam 00007420-RA
Stylophora pistillata|Hood coral|Spis14035.t1 SpisGene14035
Fungia spp.|Coral|ffun1.m4.21870.m1
Goniastrea aspera|Stony coral|gasp1.m3.6245.m1
Galaxea fascicularis|Galaxy coral|gfas1.m1.11287.m1
Homo sapiens|Human|TRPM8 CCDS33407.1
Homo sapiens|Human|TRPM2 CCDS13710.1
Homo sapiens|Human|TRPM5 CCDS31340.1
Homo sapiens|Human|TRPM4 CCDS33073.1
Acanthaster planci|Crown-of-thorns starfish|XP 022086910.1
Ptychodera flava|Acorn worm|pfl 40v0 9 20150316 1g24145.
Saccoglossus kowalevskii|Acorn worm|XP 006816420
Acanthaster planci|Crown-of-thorns starfish|XP 022096576.1
Strongylocentrotus purpuratus|Purple sea urchin|XP 011661004.1
Ptychodera flava|Acorn worm|pfl 40v0 9 20150316 1g5514.
Ptychodera flava|Acorn worm|pfl 40v0 9 20150316 1g7524.
Ptychodera flava|Acorn worm|pfl 40v0 9 20150316 1g8370.
Ptychodera flava|Acorn worm|pfl 40v0 9 20150316 1g8628.
Ptychodera flava|Acorn worm|pfl 40v0 9 20150316 1g26243.
Strongylocentrotus purpuratus|Purple sea urchin|XP 011664673.1
Apostichopus japonicus|Japanese sea cucumber|PIK58237.1
Acanthaster planci|Crown-of-thorns starfish|XP 022110675.1
Acanthaster planci|Crown-of-thorns starfish|XP 022110674.1
Strongylocentrotus purpuratus|Purple sea urchin|XP 011674872.1
Apostichopus japonicus|Japanese sea cucumber|PIK46475.1
Acanthaster planci|Crown-of-thorns starfish|XP 022096929.1
Acanthaster planci|Crown-of-thorns starfish|XP 022084590.1
Acanthaster planci|Crown-of-thorns starfish|XP 022084599.1
α
Saccoglossus kowalevskii|Acorn worm|XP 006814003
Strongylocentrotus purpuratus|Purple sea urchin|XP 794425.3
Acanthaster planci|Crown-of-thorns starfish|XP 022099673.1
Drosophila melanogaster|Common fruit fly|Trpm
Homo sapiens|Human|TRPM3 CCDS6634.1
Homo sapiens|Human|TRPM1 CCDS10024.2
Homo sapiens|Human|TRPM7 CCDS42035.1
Homo sapiens|Human|TRPM6 CCDS6647.1
Hydra vulgaris|Fresh-water polyp|XP 012565257
Aurelia|Moon jelly|Seg1831.8 transcript id Seg1831.8|GoldUCD|mRNA.D3Y31
Nematostella vectensis|Starlet sea anemone|Nemve1|248535
Porites lutea|Small polyp stony coral|plut2.m8.2561.m1
Galaxea fascicularis|Galaxy coral|gfas1.m1.6599.m1
Acropora digitifera|Coral|adi v1.10270
Acropora tenuis|Coral|aten 0.1.m1.1851.m1
Fungia spp.|Coral|ffun1.m4.19256.m1
Goniastrea aspera|Stony coral|gasp1.m3.30302.m1
Pocillopora damicornis|Lace coral|pdam 00009856-RA
Stylophora pistillata|Hood coral|Spis4675 SpisGene4675
Hydra vulgaris|Fresh-water polyp|XP 012554388
Hydra vulgaris|Fresh-water polyp|XP 012562662
Hydra vulgaris|Fresh-water polyp|XP 012561335
Aiptasia|Sea anemone|AIPGENE12924
Nematostella vectensis|Starlet sea anemone|Nemve1|85952
Pocillopora damicornis|Lace coral|pdam 00014050-RA
Goniastrea aspera|Stony coral|gasp1.m3.26670.m1
Fungia spp.|Coral|ffun1.m4.18855.m1
Aiptasia|Sea anemone|AIPGENE19094
Nematostella vectensis|Starlet sea anemone|Nemve1|102167
Porites lutea|Small polyp stony coral|plut2.m8.21682.m1
Pocillopora damicornis|Lace coral|pdam 00020220-RA
Goniastrea aspera|Stony coral|gasp1.m3.6037.m1
Fungia spp.|Coral|ffun1.m4.18290.m1
Duplications before
Cniadaria-Bilateria split

Figure S12

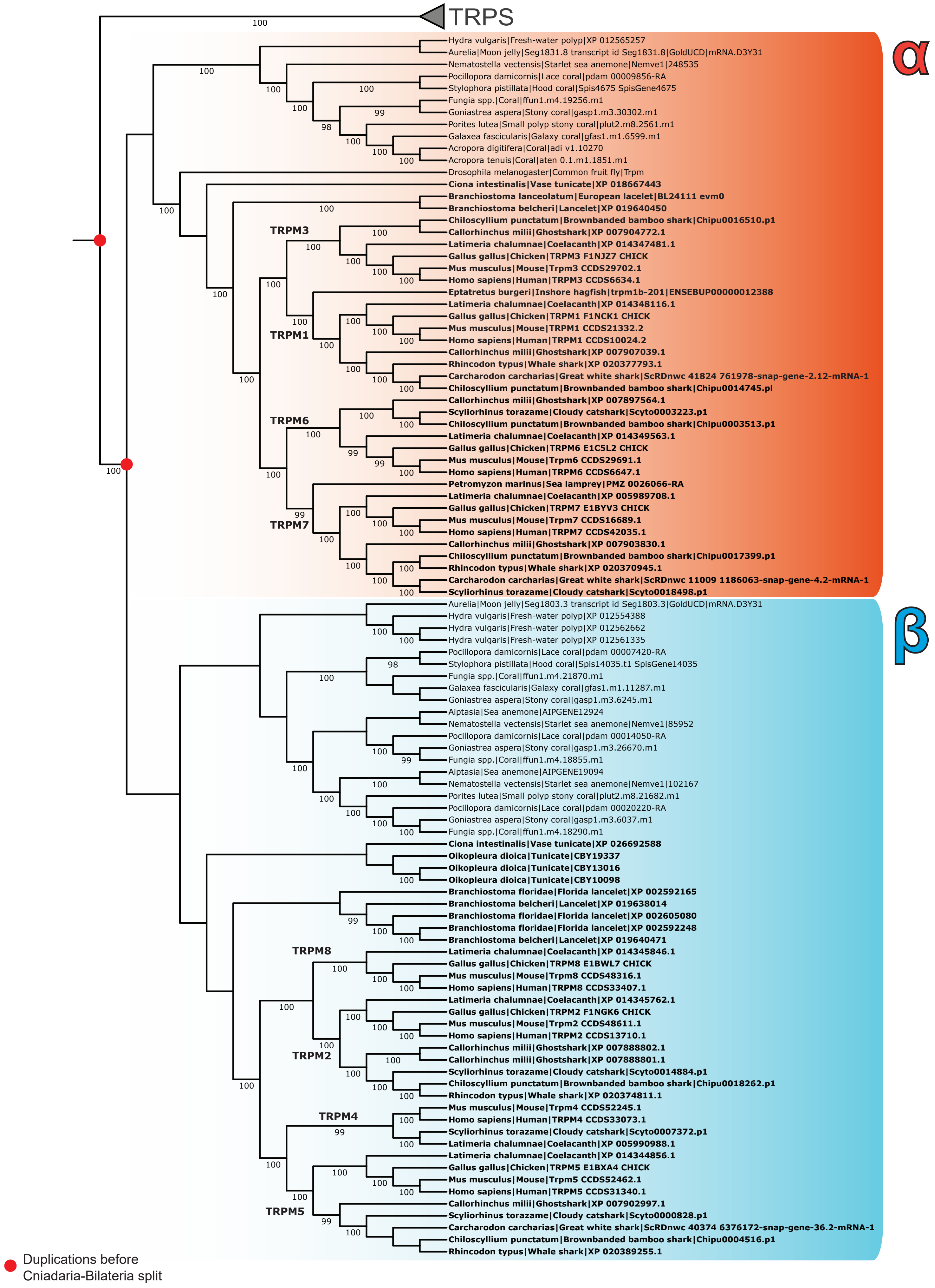
TRPS
α
Hydra vulgaris|Fresh-water polyp|XP 012565257
Aurelia|Moon jelly|Seg1831.8 transcript id Seg1831.8|GoldUCD|mRNA.D3Y31
Nematostella vectensis|Starlet sea anemone|Nemve1|248535
Pocillopora damicornis|Lace coral|pdam 00009856-RA
Stylophora pistillata|Hood coral|Spis4675 SpisGene4675
Fungia spp.|Coral|ffun1.m4.19256.m1
Goniastrea aspera|Stony coral|gasp1.m3.30302.m1
Porites lutea|Small polyp stony coral|plut2.m8.2561.m1
Galaxea fascicularis|Galaxy coral|gfas1.m1.6599.m1
Acropora digitifera|Coral|adi v1.10270
Acropora tenuis|Coral|aten 0.1.m1.1851.m1
Drosophila melanogaster|Common fruit fly|Trpm
Ciona intestinalis|Vase tunicate|XP 018667443
Branchiostoma lanceolatum|European lacelet|BL24111 evm0
Branchiostoma belcheri|Lancelet|XP 019640450
TRPM3
Chiloscyllium punctatum|Brownbanded bamboo shark|Chipu0016510.p1
Callorhinchus milii|Ghostshark|XP 007904772.1
Latimeria chalumnae|Coelacanth|XP 014347481.1
Gallus gallus|Chicken|TRPM3 F1NJZ7 CHICK
Mus musculus|Mouse|Trpm3 CCDS29702.1
Homo sapiens|Human|TRPM3 CCDS6634.1
Eptatretus burgeri|Inshore hagfish|trpm1b-201|ENSEBUP00000012388
TRPM1
Latimeria chalumnae|Coelacanth|XP 014348116.1
Gallus gallus|Chicken|TRPM1 F1NCK1 CHICK
Mus musculus|Mouse|TRPM1 CCDS21332.2
Homo sapiens|Human|TRPM1 CCDS10024.2
Callorhinchus milii|Ghostshark|XP 007907039.1
Rhincodon typus|Whale shark|XP 020377793.1
Carcharodon carcharias|Great white shark|ScRDnwc 41824 761978-snap-gene-2.12-mRNA-1
Chiloscyllium punctatum|Brownbanded bamboo shark|Chipu0014745.pl
TRPM6
Callorhinchus milii|Ghostshark|XP 007897564.1
Scyliorhinus torazame|Cloudy catshark|Scyto0003223.p1
Chiloscyllium punctatum|Brownbanded bamboo shark|Chipu0003513.p1
Latimeria chalumnae|Coelacanth|XP 014349563.1
Gallus gallus|Chicken|TRPM6 E1C5L2 CHICK
Mus musculus|Mouse|Trpm6 CCDS29691.1
Homo sapiens|Human|TRPM6 CCDS6647.1
Petromyzon marinus|Sea lamprey|PMZ 0026066-RA
TRPM7
Latimeria chalumnae|Coelacanth|XP 005989708.1
Gallus gallus|Chicken|TRPM7 E1BYV3 CHICK
Mus musculus|Mouse|Trpm7 CCDS16689.1
Homo sapiens|Human|TRPM7 CCDS42035.1
Callorhinchus milii|Ghostshark|XP 007903830.1
Chiloscyllium punctatum|Brownbanded bamboo shark|Chipu0017399.p1
Rhincodon typus|Whale shark|XP 020370945.1
Carcharodon carcharias|Great white shark|ScRDnwc 11009 1186063-snap-gene-4.2-mRNA-1
Scyliorhinus torazame|Cloudy catshark|Scyto0018498.p1
β
Aurelia|Moon jelly|Seg1803.3 transcript id Seg1803.3|GoldUCD|mRNA.D3Y31
Hydra vulgaris|Fresh-water polyp|XP 012554388
Hydra vulgaris|Fresh-water polyp|XP 012562662
Hydra vulgaris|Fresh-water polyp|XP 012561335
Pocillopora damicornis|Lace coral|pdam 00007420-RA
Stylophora pistillata|Hood coral|Spis14035.t1 SpisGene14035
Fungia spp.|Coral|ffun1.m4.21870.m1
Galaxea fascicularis|Galaxy coral|gfas1.m1.11287.m1
Goniastrea aspera|Stony coral|gasp1.m3.6245.m1
Aiptasia|Sea anemone|AIPGENE12924
Nematostella vectensis|Starlet sea anemone|Nemve1|85952
Pocillopora damicornis|Lace coral|pdam 00014050-RA
Goniastrea aspera|Stony coral|gasp1.m3.26670.m1
Fungia spp.|Coral|ffun1.m4.18855.m1
Aiptasia|Sea anemone|AIPGENE19094
Nematostella vectensis|Starlet sea anemone|Nemve1|102167
Porites lutea|Small polyp stony coral|plut2.m8.21682.m1
Pocillopora damicornis|Lace coral|pdam 00020220-RA
Goniastrea aspera|Stony coral|gasp1.m3.6037.m1
Fungia spp.|Coral|ffun1.m4.18290.m1
Ciona intestinalis|Vase tunicate|XP 026692588
Oikopleura dioica|Tunicate|CBY19337
Oikopleura dioica|Tunicate|CBY13016
Oikopleura dioica|Tunicate|CBY10098
Branchiostoma floridae|Florida lancelet|XP 002592165
Branchiostoma belcheri|Lancelet|XP 019638014
Branchiostoma floridae|Florida lancelet|XP 002605080
Branchiostoma floridae|Florida lancelet|XP 002592248
Branchiostoma belcheri|Lancelet|XP 019640471
TRPM8
Latimeria chalumnae|Coelacanth|XP 014345846.1
Gallus gallus|Chicken|TRPM8 E1BWL7 CHICK
Mus musculus|Mouse|Trpm8 CCDS48316.1
Homo sapiens|Human|TRPM8 CCDS33407.1
Latimeria chalumnae|Coelacanth|XP 014345762.1
Gallus gallus|Chicken|TRPM2 F1NGK6 CHICK
Mus musculus|Mouse|Trpm2 CCDS48611.1
Homo sapiens|Human|TRPM2 CCDS13710.1
TRPM2
Callorhinchus milii|Ghostshark|XP 007888802.1
Callorhinchus milii|Ghostshark|XP 007888801.1
Scyliorhinus torazame|Cloudy catshark|Scyto0014884.p1
Chiloscyllium punctatum|Brownbanded bamboo shark|Chipu0018262.p1
Rhincodon typus|Whale shark|XP 020374811.1
TRPM4
Mus musculus|Mouse|Trpm4 CCDS52245.1
Homo sapiens|Human|TRPM4 CCDS33073.1
Scyliorhinus torazame|Cloudy catshark|Scyto0007372.p1
Latimeria chalumnae|Coelacanth|XP 005990988.1
Latimeria chalumnae|Coelacanth|XP 014344856.1
Gallus gallus|Chicken|TRPM5 E1BXA4 CHICK
Mus musculus|Mouse|Trpm5 CCDS52462.1
Homo sapiens|Human|TRPM5 CCDS31340.1
TRPM5
Callorhinchus milii|Ghostshark|XP 007902997.1
Scyliorhinus torazame|Cloudy catshark|Scyto0000828.p1
Carcharodon carcharias|Great white shark|ScRDnwc 40374 6376172-snap-gene-36.2-mRNA-1
Chiloscyllium punctatum|Brownbanded bamboo shark|Chipu0004516.p1
Rhincodon typus|Whale shark|XP 020389255.1
Duplications before
Cniadaria-Bilateria split

Figure S13

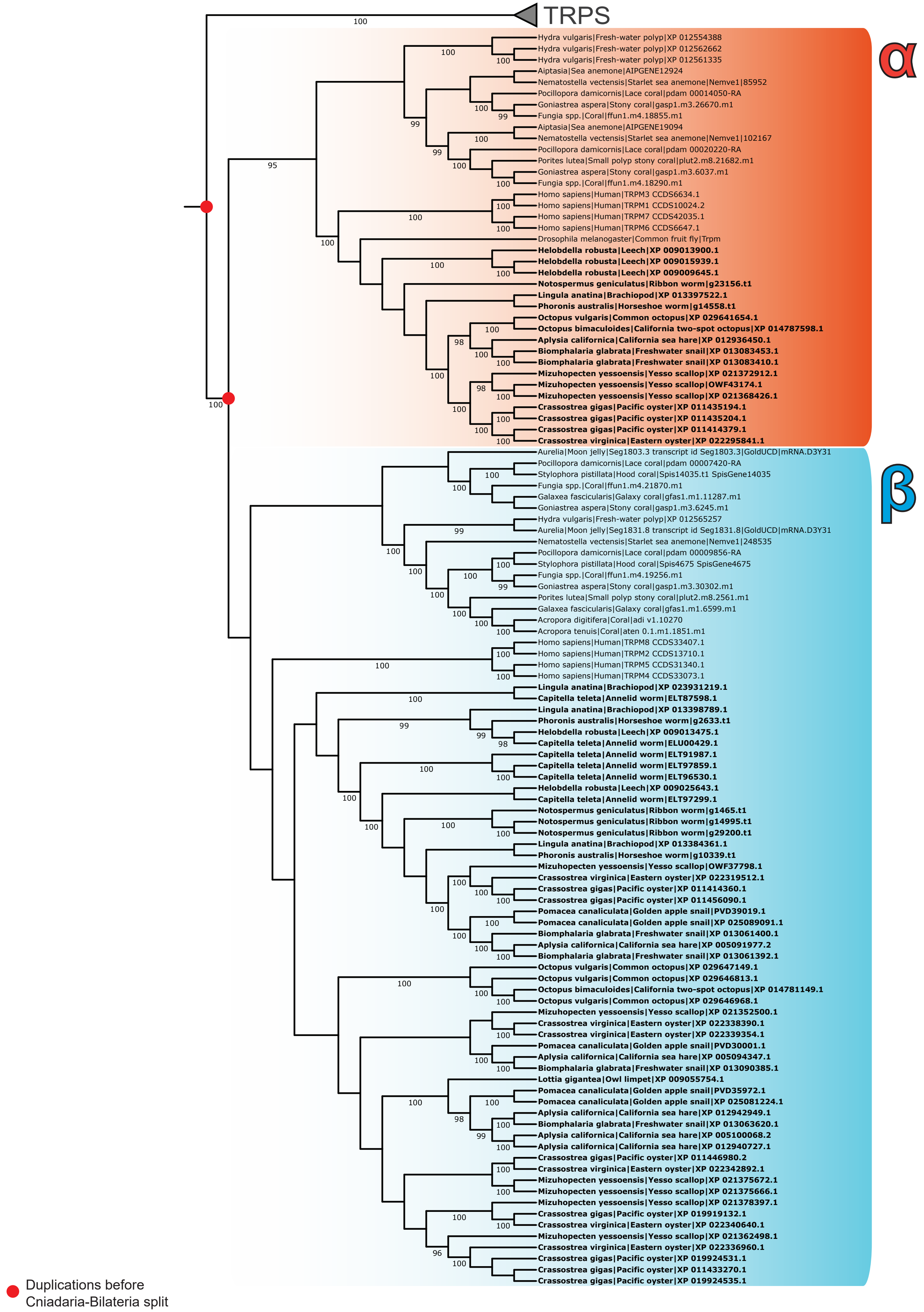

TRPS
α
Hydra vulgaris|Fresh-water polyp|XP 012554388
Hydra vulgaris|Fresh-water polyp|XP 012562662
Hydra vulgaris|Fresh-water polyp|XP 012561335
Aiptasia|Sea anemone|AIPGENE12924
Nematostella vectensis|Starlet sea anemone|Nemve1|85952
Pocillopora damicornis|Lace coral|pdam 00014050-RA
Goniastrea aspera|Stony coral|gasp1.m3.26670.m1
Fungia spp.|Coral|ffun1.m4.18855.m1
Aiptasia|Sea anemone|AIPGENE19094
Nematostella vectensis|Starlet sea anemone|Nemve1|102167
Pocillopora damicornis|Lace coral|pdam 00020220-RA
Porites lutea|Small polyp stony coral|plut2.m8.21682.m1
Goniastrea aspera|Stony coral|gasp1.m3.6037.m1
Fungia spp.|Coral|ffun1.m4.18290.m1
Homo sapiens|Human|TRPM3 CCDS6634.1
Homo sapiens|Human|TRPM1 CCDS10024.2
Homo sapiens|Human|TRPM7 CCDS42035.1
Homo sapiens|Human|TRPM6 CCDS6647.1
Drosophila melanogaster|Common fruit fly|Trpm
Helobdella robusta|Leech|XP 009013900.1
Helobdella robusta|Leech|XP 009015939.1
Helobdella robusta|Leech|XP 009009645.1
Notospermus geniculatus|Ribbon worm|g23156.t1
Lingula anatina|Brachiopod|XP 013397522.1
Phoronis australis|Horseshoe worm|g14558.t1
Octopus vulgaris|Common octopus|XP 029641654.1
Octopus bimaculoides|California two-spot octopus|XP 014787598.1
Aplysia californica|California sea hare|XP 012936450.1
Biomphalaria glabrata|Freshwater snail|XP 013083453.1
Biomphalaria glabrata|Freshwater snail|XP 013083410.1
Mizuhopecten yessoensis|Yesso scallop|XP 021372912.1
Mizuhopecten yessoensis|Yesso scallop|OWF43174.1
Mizuhopecten yessoensis|Yesso scallop|XP 021368426.1
Crassostrea gigas|Pacific oyster|XP 011435194.1
Crassostrea gigas|Pacific oyster|XP 011435204.1
Crassostrea gigas|Pacific oyster|XP 011414379.1
Crassostrea virginica|Eastern oyster|XP 022295841.1
β
Aurelia|Moon jelly|Seg1803.3 transcript id Seg1803.3|GoldUCD|mRNA.D3Y31
Pocillopora damicornis|Lace coral|pdam 00007420-RA
Stylophora pistillata|Hood coral|Spis14035.t1 SpisGene14035
Fungia spp.|Coral|ffun1.m4.21870.m1
Galaxea fascicularis|Galaxy coral|gfas1.m1.11287.m1
Goniastrea aspera|Stony coral|gasp1.m3.6245.m1
Hydra vulgaris|Fresh-water polyp|XP 012565257
Aurelia|Moon jelly|Seg1831.8 transcript id Seg1831.8|GoldUCD|mRNA.D3Y31
Nematostella vectensis|Starlet sea anemone|Nemve1|248535
Pocillopora damicornis|Lace coral|pdam 00009856-RA
Stylophora pistillata|Hood coral|Spis4675 SpisGene4675
Fungia spp.|Coral|ffun1.m4.19256.m1
Goniastrea aspera|Stony coral|gasp1.m3.30302.m1
Porites lutea|Small polyp stony coral|plut2.m8.2561.m1
Galaxea fascicularis|Galaxy coral|gfas1.m1.6599.m1
Acropora digitifera|Coral|adi v1.10270
Acropora tenuis|Coral|aten 0.1.m1.1851.m1
Homo sapiens|Human|TRPM8 CCDS33407.1
Homo sapiens|Human|TRPM2 CCDS13710.1
Homo sapiens|Human|TRPM5 CCDS31340.1
Homo sapiens|Human|TRPM4 CCDS33073.1
Lingula anatina|Brachiopod|XP 023931219.1
Capitella teleta|Annelid worm|ELT87598.1
Lingula anatina|Brachiopod|XP 013398789.1
Phoronis australis|Horseshoe worm|g2633.t1
Helobdella robusta|Leech|XP 009013475.1
Capitella teleta|Annelid worm|ELU00429.1
Capitella teleta|Annelid worm|ELT91987.1
Capitella teleta|Annelid worm|ELT97859.1
Capitella teleta|Annelid worm|ELT96530.1
Helobdella robusta|Leech|XP 009025643.1
Capitella teleta|Annelid worm|ELT97299.1
Notospermus geniculatus|Ribbon worm|g1465.t1
Notospermus geniculatus|Ribbon worm|g14995.t1
Notospermus geniculatus|Ribbon worm|g29200.t1
Lingula anatina|Brachiopod|XP 013384361.1
Phoronis australis|Horseshoe worm|g10339.t1
Mizuhopecten yessoensis|Yesso scallop|OWF37798.1
Crassostrea virginica|Eastern oyster|XP 022319512.1
Crassostrea gigas|Pacific oyster|XP 011414360.1
Crassostrea gigas|Pacific oyster|XP 011456090.1
Pomacea canaliculata|Golden apple snail|PVD39019.1
Pomacea canaliculata|Golden apple snail|XP 025089091.1
Biomphalaria glabrata|Freshwater snail|XP 013061400.1
Aplysia californica|California sea hare|XP 005091977.2
Biomphalaria glabrata|Freshwater snail|XP 013061392.1
Octopus vulgaris|Common octopus|XP 029647149.1
Octopus vulgaris|Common octopus|XP 029646813.1
Octopus bimaculoides|California two-spot octopus|XP 014781149.1
Octopus vulgaris|Common octopus|XP 029646968.1
Mizuhopecten yessoensis|Yesso scallop|XP 021352500.1
Crassostrea virginica|Eastern oyster|XP 022338390.1
Crassostrea virginica|Eastern oyster|XP 022339354.1
Pomacea canaliculata|Golden apple snail|PVD30001.1
Aplysia californica|California sea hare|XP 005094347.1
Biomphalaria glabrata|Freshwater snail|XP 013090385.1
Lottia gigantea|Owl limpet|XP 009055754.1
Pomacea canaliculata|Golden apple snail|PVD35972.1
Pomacea canaliculata|Golden apple snail|XP 025081224.1
Aplysia californica|California sea hare|XP 012942949.1
Biomphalaria glabrata|Freshwater snail|XP 013063620.1
Aplysia californica|California sea hare|XP 005100068.2
Aplysia californica|California sea hare|XP 012940727.1
Crassostrea gigas|Pacific oyster|XP 011446980.2
Crassostrea virginica|Eastern oyster|XP 022342892.1
Mizuhopecten yessoensis|Yesso scallop|XP 021375672.1
Mizuhopecten yessoensis|Yesso scallop|XP 021375666.1
Mizuhopecten yessoensis|Yesso scallop|XP 021378397.1
Crassostrea gigas|Pacific oyster|XP 019919132.1
Crassostrea virginica|Eastern oyster|XP 022340640.1
Mizuhopecten yessoensis|Yesso scallop|XP 021362498.1
Crassostrea virginica|Eastern oyster|XP 022336960.1
Crassostrea gigas|Pacific oyster|XP 019924531.1
Crassostrea gigas|Pacific oyster|XP 011433270.1
Crassostrea gigas|Pacific oyster|XP 019924535.1
Duplications before
Cniadaria-Bilateria split


Figure S14

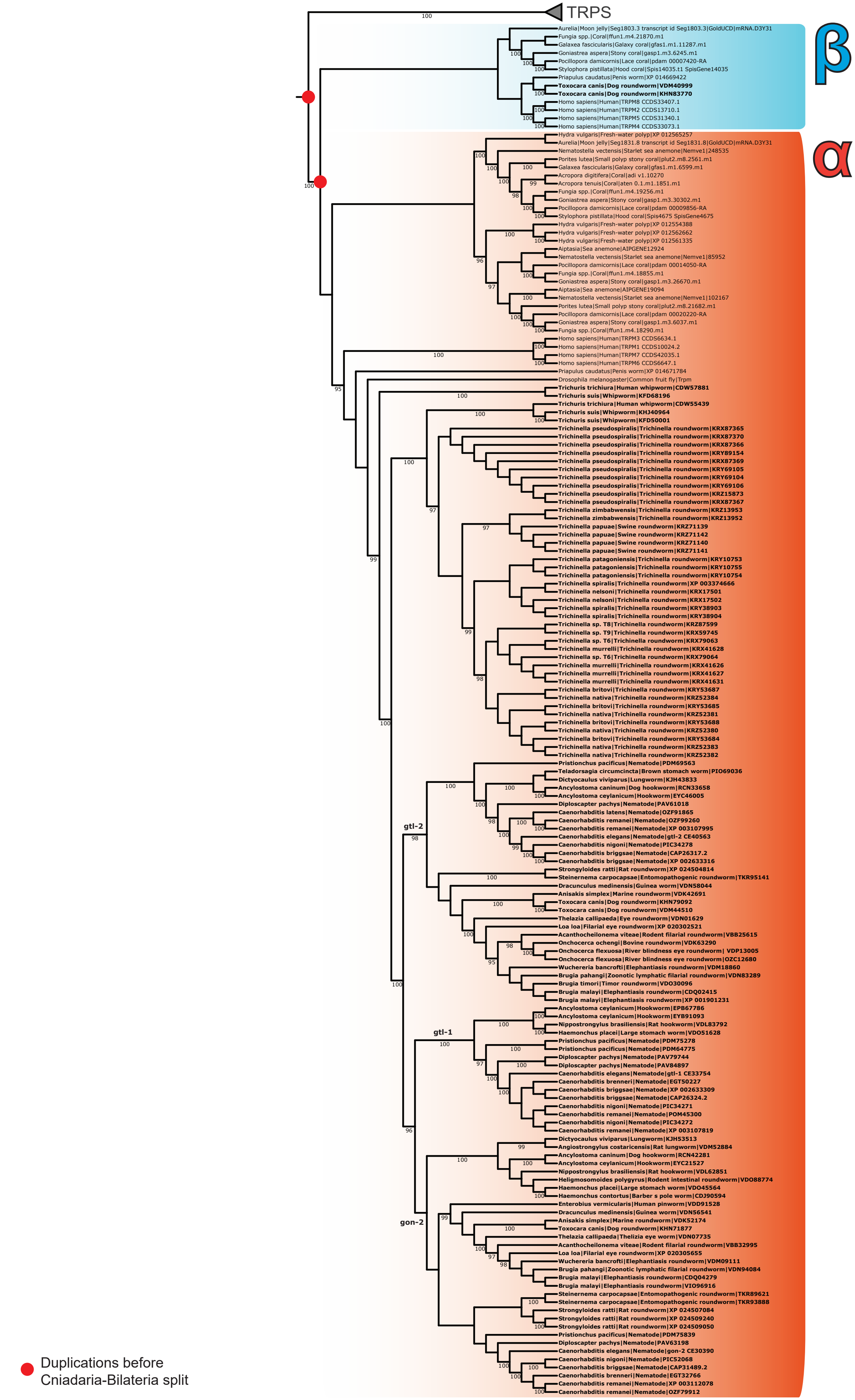
TRPS
β
α
Aurelia|Moon jelly|Seg1803.3 transcript id Seg1803.3|GoldUCD|mRNA.D3Y31
Fungia spp.|Coral|ffun1.m4.21870.m1
Galaxea fascicularis|Galaxy coral|gfas1.m1.11287.m1
Goniastrea aspera|Stony coral|gasp1.m3.6245.m1
Pocillopora damicornis|Lace coral|pdam 00007420-RA
Stylophora pistillata|Hood coral|Spis14035.t1 SpisGene14035
Priapulus caudatus|Penis worm|XP 014669422
Toxocara canis|Dog roundworm|VDM40999
Toxocara canis|Dog roundworm|KHN83770
Homo sapiens|Human|TRPM8 CCDS33407.1
Homo sapiens|Human|TRPM2 CCDS13710.1
Homo sapiens|Human|TRPM5 CCDS31340.1
Homo sapiens|Human|TRPM4 CCDS33073.1
Hydra vulgaris|Fresh-water polyp|XP 012565257
Aurelia|Moon jelly|Seg1831.8 transcript id Seg1831.8|GoldUCD|mRNA.D3Y31
Nematostella vectensis|Starlet sea anemone|Nemve1|248535
Porites lutea|Small polyp stony coral|plut2.m8.2561.m1
Galaxea fascicularis|Galaxy coral|gfas1.m1.6599.m1
Acropora digitifera|Coral|adi v1.10270
Acropora tenuis|Coral|aten 0.1.m1.1851.m1
Fungia spp.|Coral|ffun1.m4.19256.m1
Goniastrea aspera|Stony coral|gasp1.m3.30302.m1
Pocillopora damicornis|Lace coral|pdam 00009856-RA
Stylophora pistillata|Hood coral|Spis4675 SpisGene4675
Hydra vulgaris|Fresh-water polyp|XP 012554388
Hydra vulgaris|Fresh-water polyp|XP 012562662
Hydra vulgaris|Fresh-water polyp|XP 012561335
Aiptasia|Sea anemone|AIPGENE12924
Nematostella vectensis|Starlet sea anemone|Nemve1|85952
Pocillopora damicornis|Lace coral|pdam 00014050-RA
Fungia spp.|Coral|ffun1.m4.18855.m1
Goniastrea aspera|Stony coral|gasp1.m3.26670.m1
Aiptasia|Sea anemone|AIPGENE19094
Nematostella vectensis|Starlet sea anemone|Nemve1|102167
Porites lutea|Small polyp stony coral|plut2.m8.21682.m1
Pocillopora damicornis|Lace coral|pdam 00020220-RA
Goniastrea aspera|Stony coral|gasp1.m3.6037.m1
Fungia spp.|Coral|ffun1.m4.18290.m1
Homo sapiens|Human|TRPM3 CCDS6634.1
Homo sapiens|Human|TRPM1 CCDS10024.2
Homo sapiens|Human|TRPM7 CCDS42035.1
Homo sapiens|Human|TRPM6 CCDS6647.1
Priapulus caudatus|Penis worm|XP 014671784
Drosophila melanogaster|Common fruit fly|Trpm
Trichuris trichiura|Human whipworm|CDW57881
Trichuris suis|Whipworm|KFD68196
Trichuris trichiura|Human whipworm|CDW55439
Trichuris suis|Whipworm|KHJ40964
Trichuris suis|Whipworm|KFD50001
Trichinella pseudospiralis|Trichinella roundworm|KRX87365
Trichinella pseudospiralis|Trichinella roundworm|KRX87370
Trichinella pseudospiralis|Trichinella roundworm|KRX87366
Trichinella pseudospiralis|Trichinella roundworm|KRY89154
Trichinella pseudospiralis|Trichinella roundworm|KRX87369
Trichinella pseudospiralis|Trichinella roundworm|KRY69105
Trichinella pseudospiralis|Trichinella roundworm|KRY69104
Trichinella pseudospiralis|Trichinella roundworm|KRY69106
Trichinella pseudospiralis|Trichinella roundworm|KRZ15873
Trichinella pseudospiralis|Trichinella roundworm|KRX87367
Trichinella zimbabwensis|Trichinella roundworm|KRZ13953
Trichinella zimbabwensis|Trichinella roundworm|KRZ13952
Trichinella papuae|Swine roundworm|KRZ71139
Trichinella papuae|Swine roundworm|KRZ71142
Trichinella papuae|Swine roundworm|KRZ71140
Trichinella papuae|Swine roundworm|KRZ71141
Trichinella patagoniensis|Trichinella roundworm|KRY10753
Trichinella patagoniensis|Trichinella roundworm|KRY10755
Trichinella patagoniensis|Trichinella roundworm|KRY10754
Trichinella spiralis|Trichinella roundworm|XP 003374666
Trichinella nelsoni|Trichinella roundworm|KRX17501
Trichinella nelsoni|Trichinella roundworm|KRX17502
Trichinella spiralis|Trichinella roundworm|KRY38903
Trichinella spiralis|Trichinella roundworm|KRY38904
Trichinella sp. T8|Trichinella roundworm|KRZ87599
Trichinella sp. T9|Trichinella roundworm|KRX59745
Trichinella sp. T6|Trichinella roundworm|KRX79063
Trichinella murrelli|Trichinella roundworm|KRX41628
Trichinella sp. T6|Trichinella roundworm|KRX79064
Trichinella murrelli|Trichinella roundworm|KRX41626
Trichinella murrelli|Trichinella roundworm|KRX41627
Trichinella murrelli|Trichinella roundworm|KRX41631
Trichinella britovi|Trichinella roundworm|KRY53687
Trichinella nativa|Trichinella roundworm|KRZ52384
Trichinella britovi|Trichinella roundworm|KRY53685
Trichinella nativa|Trichinella roundworm|KRZ52381
Trichinella britovi|Trichinella roundworm|KRY53688
Trichinella nativa|Trichinella roundworm|KRZ52380
Trichinella britovi|Trichinella roundworm|KRY53684
Trichinella nativa|Trichinella roundworm|KRZ52383
Trichinella nativa|Trichinella roundworm|KRZ52382
Pristionchus pacificus|Nematode|PDM69563
Teladorsagia circumcincta|Brown stomach worm|PIO69036
Dictyocaulus viviparus|Lungworm|KJH43833
Ancylostoma caninum|Dog hookworm|RCN33658
Ancylostoma ceylanicum|Hookworm|EYC46005
Diploscapter pachys|Nematode|PAV61018
Caenorhabditis latens|Nematode|OZF91865
Caenorhabditis remanei|Nematode|OZF99260
Caenorhabditis remanei|Nematode|XP 003107995
Caenorhabditis elegans|Nematode|gtl-2 CE40563
Caenorhabditis nigoni|Nematode|PIC34278
Caenorhabditis briggsae|Nematode|CAP26317.2
Caenorhabditis briggsae|Nematode|XP 002633316
Strongyloides ratti|Rat roundworm|XP 024504814
Steinernema carpocapsae|Entomopathogenic roundworm|TKR95141
Dracunculus medinensis|Guinea worm|VDN58044
Anisakis simplex|Marine roundworm|VDK42691
Toxocara canis|Dog roundworm|KHN79092
Toxocara canis|Dog roundworm|VDM44510
Thelazia callipaeda|Eye roundworm|VDN01629
Loa loa|Filarial eye roundworm|XP 020302521
Acanthocheilonema viteae|Rodent filarial roundworm|VBB25615
Onchocerca ochengi|Bovine roundworm|VDK63290
Onchocerca flexuosa|River blindness eye roundworm| VDP13005
Onchocerca flexuosa|River blindness eye roundworm|OZC12680
Wuchereria bancrofti|Elephantiasis roundworm|VDM18860
Brugia pahangi|Zoonotic lymphatic filarial roundworm|VDN83289
Brugia timori|Timor roundworm|VDO30096
Brugia malayi|Elephantiasis roundworm|CDQ02415
Brugia malayi|Elephantiasis roundworm|XP 001901231
Ancylostoma ceylanicum|Hookworm|EPB67786
Ancylostoma ceylanicum|Hookworm|EYB91093
Nippostrongylus brasiliensis|Rat hookworm|VDL83792
Haemonchus placei|Large stomach worm|VDO51628
Pristionchus pacificus|Nematode|PDM75278
Pristionchus pacificus|Nematode|PDM64775
Diploscapter pachys|Nematode|PAV79744
Diploscapter pachys|Nematode|PAV84897
Caenorhabditis elegans|Nematode|gtl-1 CE33754
Caenorhabditis brenneri|Nematode|EGT50227
Caenorhabditis briggsae|Nematode|XP 002633309
Caenorhabditis briggsae|Nematode|CAP26324.2
Caenorhabditis nigoni|Nematode|PIC34271
Caenorhabditis remanei|Nematode|POM45300
Caenorhabditis nigoni|Nematode|PIC34272
Caenorhabditis remanei|Nematode|XP 003107819
Dictyocaulus viviparus|Lungworm|KJH53513
Angiostrongylus costaricensis|Rat lungworm|VDM52884
Ancylostoma caninum|Dog hookworm|RCN42281
Ancylostoma ceylanicum|Hookworm|EYC21527
Nippostrongylus brasiliensis|Rat hookworm|VDL62851
Heligmosomoides polygyrus|Rodent intestinal roundworm|VDO88774
Haemonchus placei|Large stomach worm|VDO45564
Haemonchus contortus|Barber s pole worm|CDJ90594
Enterobius vermicularis|Human pinworm|VDD91528
Dracunculus medinensis|Guinea worm|VDN56541
Anisakis simplex|Marine roundworm|VDK52174
Toxocara canis|Dog roundworm|KHN71877
Thelazia callipaeda|Thelizia eye worm|VDN07735
Acanthocheilonema viteae|Rodent filarial roundworm|VBB32995
Loa loa|Filarial eye roundworm|XP 020305655
Wuchereria bancrofti|Elephantiasis roundworm|VDM09111
Brugia pahangi|Zoonotic lymphatic filarial roundworm|VDN94084
Brugia malayi|Elephantiasis roundworm|CDQ04279
Brugia malayi|Elephantiasis roundworm|VIO96916
Steinernema carpocapsae|Entomopathogenic roundworm|TKR89621
Steinernema carpocapsae|Entomopathogenic roundworm|TKR93888
Strongyloides ratti|Rat roundworm|XP 024507084
Strongyloides ratti|Rat roundworm|XP 024509240
Strongyloides ratti|Rat roundworm|XP 024509050
Pristionchus pacificus|Nematode|PDM75839
Diploscapter pachys|Nematode|PAV63198
Caenorhabditis elegans|Nematode|gon-2 CE30390
Caenorhabditis nigoni|Nematode|PIC52068
Caenorhabditis briggsae|Nematode|CAP31489.2
Caenorhabditis brenneri|Nematode|EGT32766
Caenorhabditis remanei|Nematode|XP 003112078
Caenorhabditis remanei|Nematode|OZF79912
gtl-2
gtl-1
gon-2
Duplications before
Cniadaria-Bilateria split

Figure S15

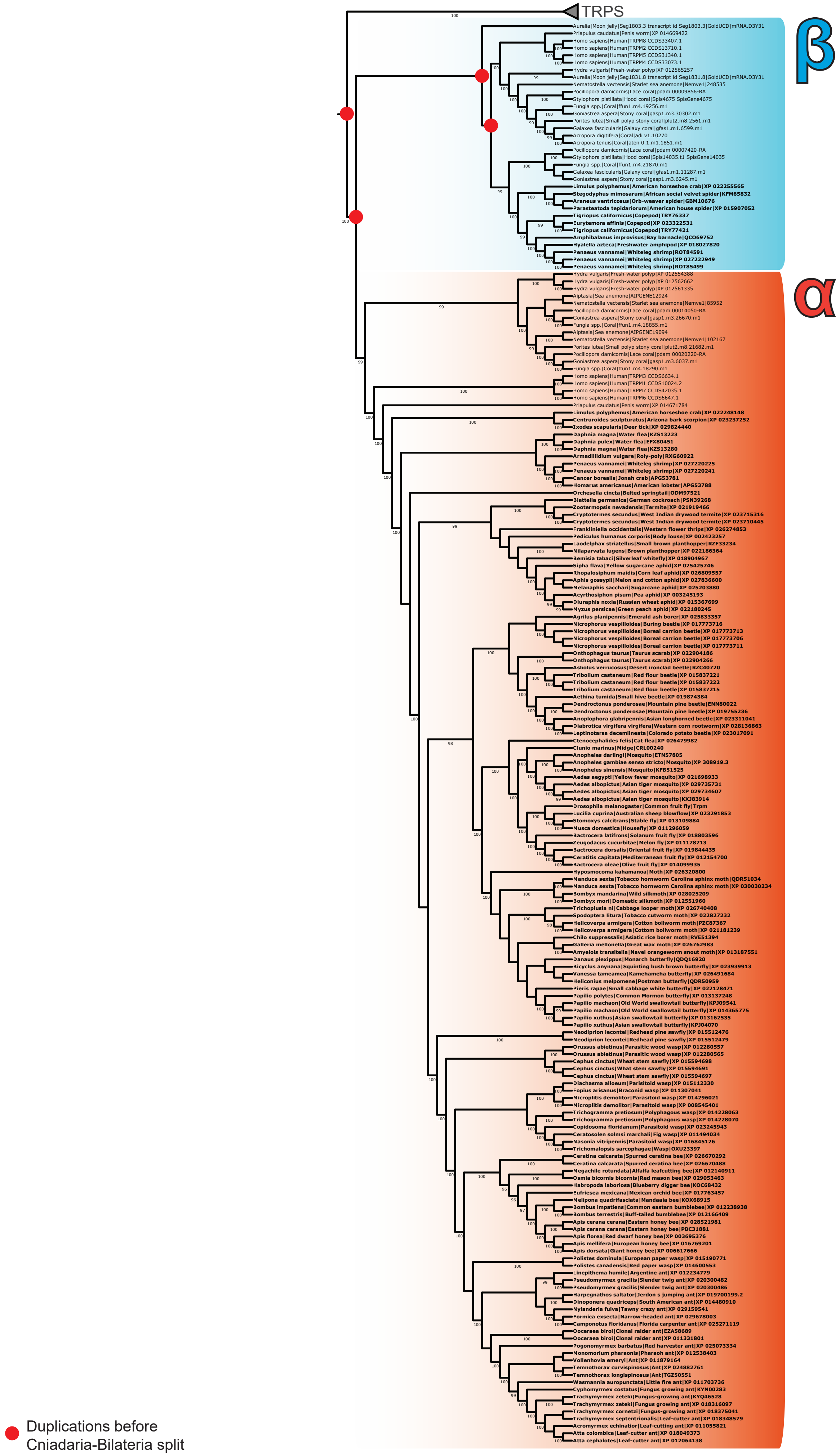


Figure S16

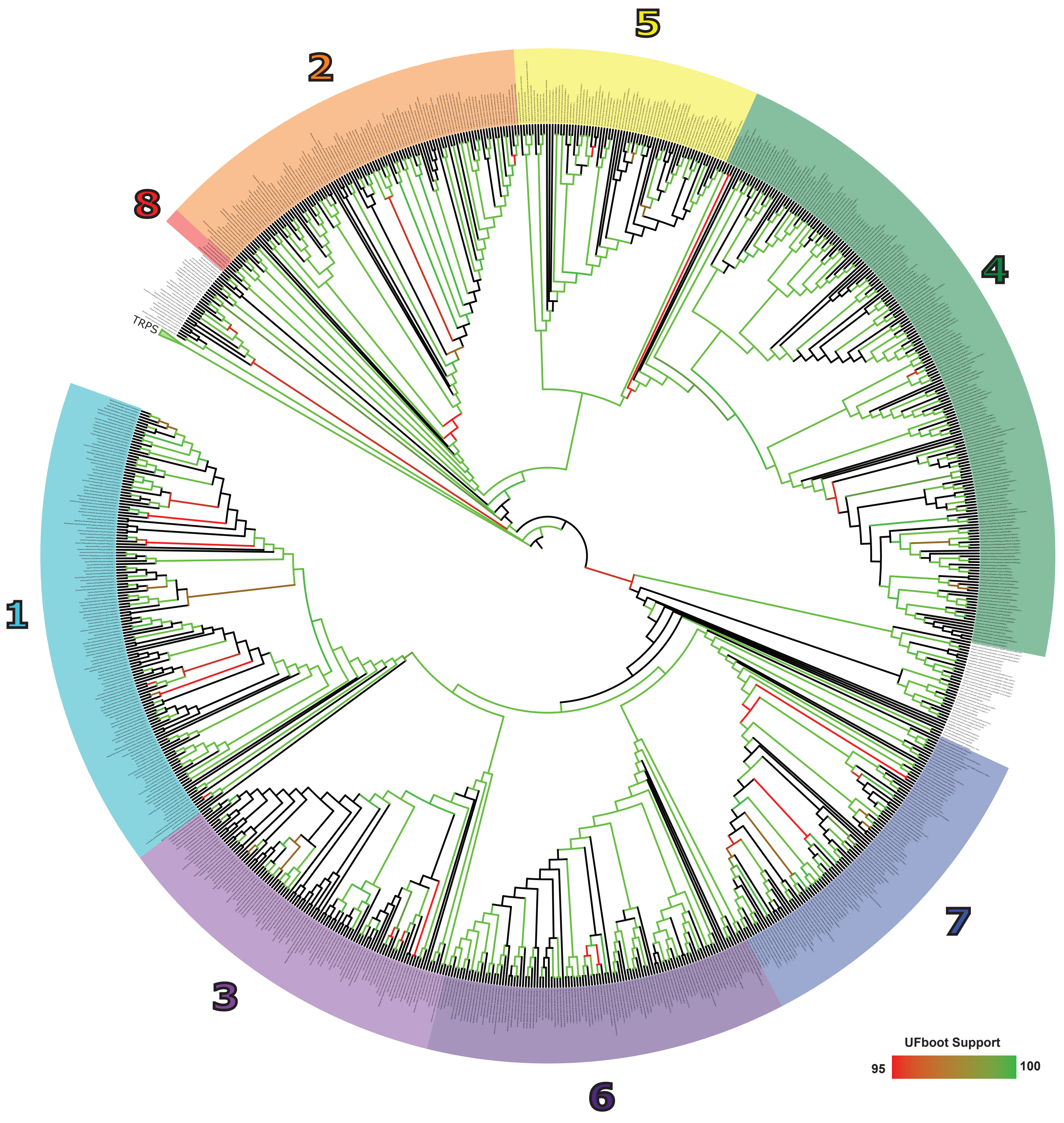


Figure S17
